# Sialidases and Fucosidases of *Akkermansia muciniphila* are crucial for growth on mucin and nutrient sharing with mucus-associated gut bacteria

**DOI:** 10.1101/2022.09.10.507281

**Authors:** Bashar Shuoker, Michael J. Pichler, Chunsheng Jin, Sakanaka Hiroka, Haiyang Wu, Ana Martínez Gascueña, Jining Liu, Tine Sofie Nielsen, Jan Holgersson, Eva Nordberg Karlsson, Nathalie Juge, Sebastian Meier, Jens Preben Morth, Niclas G. Karlsson, Maher Abou Hachem

**Affiliations:** Department of Biotechnology and Biomedicine, Technical University of Denmark, Lyngby, 2800, Denmark; Biotechnology, Department of Chemistry, Lund University, Lund, Sweden; Proteomics Core Facility at Sahlgrenska Academy, University of Gothenburg, Gothenburg, Sweden; Quadram Institute Bioscience, Norwich, United Kingdom; Department of Laboratory Medicine, Institute of Biomedicine, Sahlgrenska Academy, University of Gothenburg, Gothenburg, Sweden; Department of Chemistry, Technical University of Denmark, Kgs Lyngby, Denmark

## Abstract

The gut mucolytic specialist *Akkermansia muciniphila* is strongly associated with the integrity of the mucus layer. Mucin glycan utilization requires the removal of diverse protective caps, notably, fucose and sialic acid, but the enzymatic details of this process remain largely unknown. Here, we describe the specificities of ten *A. muciniphila* glycoside hydrolases, which collectively remove all known sialyl and fucosyl mucin caps including those with double sulphated epitopes. Structural analyses revealed an unprecedented fucosidase modular arrangement and explained the exclusive sialyl T-antigen specificity of a sialidase of a previously unknown family and catalytic apparatus. Key cell attached sialidases and fucosidases conferred mucin-binding and their inhibition abolished growth of *A. muciniphila* on mucin. Remarkably, the sialic acid fucose did not contribute to *A. muciniphila* growth, but instead promoted butyrate production by co-cultured Clostridia. This study brings unique mechanistic insight into the initiation of mucin *O*-glycan degradation by *A. muciniphila* and the nutrient sharing between key mucus-associated bacteria.

## Introduction

The gut microbiota (GM) exerts a major impact on the host immune and metabolic homeostasis^1, 2^. Fiber intake promotes a balanced microbiota, whereas a fiber-poor diet is associated with aberrant GM signatures and breach of the gut barrier^3, 4^. The host’s first defense line against microbial insult is the intestinal mucosal barrier that shields epithelial cells from the GM and increases in thickness towards the colon^5, 6^. Mucins are the main structural and gel-forming scaffolds of the mucosa, which is dominated by Mucin 2 (Muc2) in the colon^7^. Muc2, similarly to other mucins, is an *O*-glycoprotein that is secreted by intestinal goblet cells and consist of up to 80% (w/w) glycans^8^.

The colonization of mucus and its resistance to proteolytic degradation are attributed to the diversity of the mucin *O*-glycans (>100 structures reported)^9^. These glycan epitopes exhibit longitudinal variations along the gastrointestinal tract (GIT)^10^, thereby creating variable adhesion and nutritional niches for the microbiota. In humans, the *O*-glycans in the small intestine and cecum regions are densely fucosylated, with a decreasing gradient toward the distal colon, whereas an increasing gradient of sialylation and sulphation^9, 10^ is observed. A reverse gradient is observed in mice^11^. Recently, a single extracellular sulphatase was shown to be critical for the growth of *Bacteroides* spp. on densely-sulphated mucin^12^. Similarly, presence of a specific sialidase is critical to growth of *Ruminococcus gnavus* on mucin^13^.

Only a few GM members can grow on mucin as a sole carbon source^5, 14^. Atypically, *Akkermansia muciniphila* relies solely on mucin and related host-derived glycans for growth^15^, which is reflected by its large carbohydrate active enzyme (CAZyme) repertoire^16^. *A. muciniphila* has received extensive attention due to its relative abundance in healthy hosts, as opposed to patients of gut inflammatory bowel disease, including Crohn’s disease and ulcerative colitis (UC)^17^, and obesity^18^. Notably, *A. muciniphila*, positive association with Parkinson disease was also reported^19^, suggesting complex interactions with the mucus-associated GM, manifested beyond the gut niche^20^. Remarkably, the molecular details of mucin *O*-glycan deconstruction by *A. muciniphila* exoglycosidases, and especially the sialic acid and fucose decapping apparatus, remain largely an uncharted territory.

Here, we used a panel of mucins to characterize the enzymes that collectively grant *A. muciniphila* access to all known fucosyl- and sialyl-mucin epitopes. Biochemical and structural and microbiological studies allowed us to identify the key fucosidases and sialidases and. We also investigated the contribution of the characterized enzyme panel to growth on mucin and on sharing mucin-derived saccharides with Firmicutes members that are known to share the same biogeography with *A. muciniphila*. Our findings promote a mechanistic understanding of initial steps of mucin turnover by *A. muciniphila* and the importance of its unselfish mucin utilization strategy in supporting other members of the mucus-adherent GM community.

## Results

### A. muciniphila encodes six divergent fucosidases

The genome of *A. muciniphila* encodes six fucosidases, four assigned into glycoside hydrolase family 29 (GH29) and two into GH95 in the CAZy database^21^. These enzymes are henceforth designated as *Am*GH29A (locus tag Amuc_0010), *Am*GH29B (Amuc_0146) *Am*GH29C (Amuc_0392), *Am*GH29D (Amuc_0846), *Am*GH95A (Amuc_0186) and *Am*GH95B (Amuc_1120) (Supplementary Fig. 1a). All enzymes possess signal peptides, indicating non-cytoplasmic localization (Supplementary Table 1). The GH29-enzymes have variable architectures, with *Am*GH29C and *Am*GH29D being the most complex and sharing two putative carbohydrate binding modules (CBMs) (Supplementary Fig. 1a, Supplementary Fig. 2). The fucosidase catalytic modules exhibit high sequence diversity (Supplementary Fig 1b) and populate hitherto undescribed clusters in the phylogenetic trees of GH29 and GH95 sequences in CAZy (Supplementary Fig. 3).

### Two key enzymes responsible for the defucosylation of mucin and structurally related glycans

We produced and purified all six enzymes and determined their kinetic parameters towards *para*-nitrophenyl-α-L-fucoside (*p*NPFuc) (Supplementary Table 2).

Next, we assayed the enzymes against a panel of fuco-oligosaccharides (Supplementary Fig. 4). The main activity of *Am*GH29A and *Am*GH29B was on the Fucα1,3GlcNAc disaccharide (Supplementary Fig. 4a). By contrast, *Am*GH29C and *Am*GH29D were active on larger oligosaccharides from mothers’ milk (HMOs) or Lewis (Le) epitopes (Supplementary Fig. 4a-d, f-j). Both *Am*GH29C and *Am*GH29D hydrolyze 3-fucosyl lactose (3FL), but only *Am*GH29C can access this motif in the extended structure LNFP V (Supplementary Fig. 4f,k,m). However, the profiles of these two enzymes towards α-1,4-fucosyl in Le^a/b^ motifs were similar (Supplementary Fig. 4j-l). Key differences between *Am*GH29C and *Am*GH29D were the activity of *Am*GH29C but not *Am*GH29D towards the sialyl Le^a^ tetrasaccharide (Supplementary Fig 4n) and the activity of *Am*GH29D towards 2’FL (Supplementary Fig. 4e).

The two GH95 enzymes were active on Fucα1,2Gal and 2’FL (Supplementary Fig. 4c,e). Only *Am*GH95A exhibited activity on Fucα1,3GlcNAc, but not on the galactosyl-extended Le^x^ epitope (Supplementary Fig. 4a,h). By contrast, only *Am*GH95B hydrolyzed α1,2Fuc linkages in the Le^b^ tetra- and hexasaccharide epitopes (Supplementary Fig. 4i,j) and additionally hydrolyzed α1,3Fuc linkages in 3FL (Supplementary Fig. 4f). These data indicate that *Am*GH95B possesses a broader substrate range than *Am*GH95A.

Hitherto reported regioselectivities in GH29^22^ and GH95^23, 24^ stem from measurements on simple model oligosaccharides. The association of *A. muciniphila* with the mucin niche prompted us to evaluate fucosidase specificities on complex mucin *O*-glycans.

We used a mixture of purified porcine gastric mucin (PGM), porcine colonic mucin (PCM) and bovine fetuin (BF). Despite differences as compared to human intestinal mucins, this substrate combination is powerful due to the large (160 structures) *O*-glycan diversity (Supplementary Table 3) and the presence of dense sulphation and sialylation in PCM similarly to the human counterpart. We evaluated activity on blood group A, H types 1-3 and four Le epitopes (Fig. 1a,b). No reliable activity of *Am*GH29A and *Am*GH29B was observed on the analyzed glycans. By contrast, *Am*GH29C and *Am*GH29D were active on all Le, but not H-epitopes (Fig. 1a,b). Both enzymes could accommodate single fucosylated Le^x^ and Le^a^ motifs, as well as double fucosylation (Le^b/y^). (Fig. 1a,b; Supplementary Fig. 5a-c). *Am*GH29D displayed a lower overall de-fucosylation yield than *Am*GH29C on single (Le^x^), double (Le^y^) or bifurcated fucosylated-epitopes, as well as core 1-4 structures (Fig. 1a,b, Supplementary Fig. 5a-f). The lack of activity of *Am*GH29D on internal epitopes is consistent with its observed lower efficacy (Fig. 1c).

**Fig. 1:**
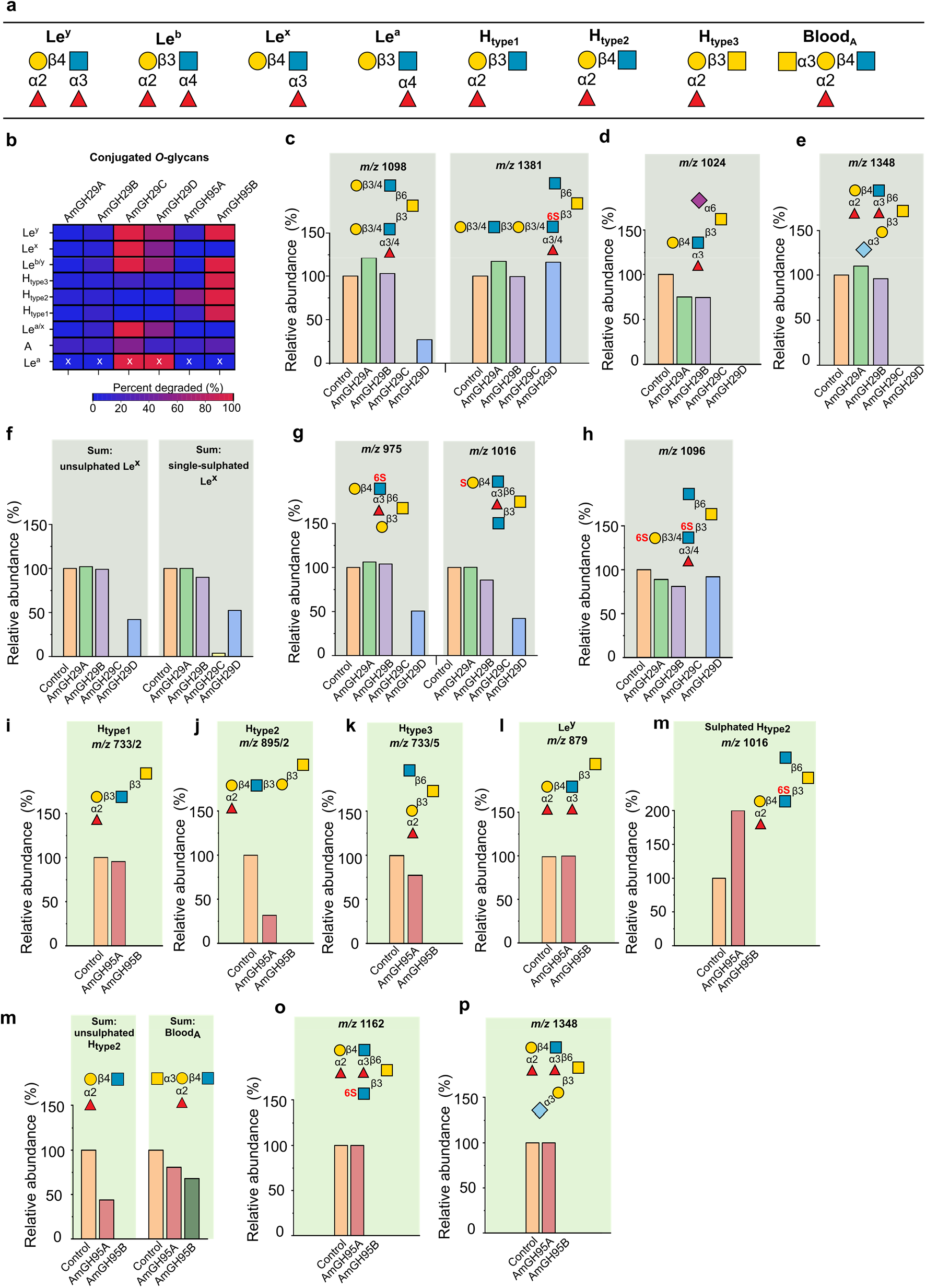
The activity profiles of the *A. muciniphila* fucosidases on mucin *O*-glycans: **a**, Overview of the fucosylated epitopes present in the analyzed conjugated *O*-glycans from intact porcine gastric and colonic mucins as well as fetuin. **b**, The fucosidase activity heat map on different epitopes. **c**, Activity profiles that reveal the sensitivity of the activity toward the fucosylation position in the glycan chain. **d-h**, Selected examples illustrating the impact of sulphation (denoted with a red capital “S”) and sialyl substitutions on fucosidase activity. **i-p**, Selected examples that demonstrate the differences in the activity profiles of the α1,2-fucosidases. Data are from a single experiment. The “x” marked data are obtained from a single glycan structure, attributed to the low abundance of assigned Le^a^ epitope based on the MS data. The linkage of the sulphatyl substituent is left out, when the assignment is not possible based on the MS data. For isobaric glycans structures with identical *m*/*z*, the additional number after the slash (/n) denotes the corresponding structure in the LC-MS data file (see extended data).

Both *Am*GH29C and *Am*GH29D were active on Le epitopes with a sialylated adjacent branch (Fig. 1d,e). In addition, a single sulphation at either the Gal or GlcNAc of Le^x^ epitopes is tolerated (Fig. 1f,g), indicating that mono-sulphation is not restricting de-fucosylation. Notably, only *Am*GH29C, but not *Am*GH29D, showed activity on the double sulphated terminal Le^x^ epitope (Fig 1h) highlighting the overall broader epitope specificity of *Am*GH29C.

*Am*GH95A and *Am*GH95B share the α1,2-fucosidases activity (Supplementary Fig. 5g-j). However, marked differences in their epitope specificity and tolerance to non-fucosyl substitutions were observed (Fig. 1b). *Am*GH95A is specific for the H2 but lacks activity on H1 and H3 epitopes (Fig. 1i-k). The activity of *Am*GH95A is also impaired by double fucosylation, *e*.*g*. in Le^y^ (Fig. 1l) and sulphation (Fig. 1m). The exclusive H type 2-specificity of *Am*GH95A is unprecedented amongst hitherto described fucosidases.

By contrast, *Am*GH95B has high activity on all H type epitopes, double fucosylated Le^y^ structures, sulphated H2 epitopes (Fig. 1i-n), and Le^y^ epitopes at sulphated (Fig. 1o) or sialylated branches at core structures (Fig. 1p). Both enzymes were sensitive to non-reducing end substitution of the H-antigen (Fig. 1n) and none were active on Fucα1,6-linked core of *N*-glycans (Supplementary Fig. 5k).

To unambiguously confirm enzyme regioselectivities on mucin-type glycoproteins, we harnessed glyco-engineered CHO cells, which display defined Lewis epitopes. *Am*GH29C and *Am*GH29D showed activity on conjugated Fucα1,3/4 linkages in Le^a/x^ and Le^b/y^ (Supplementary Fig. 6a-d), while *Am*GH95B hydrolyzed Fucα1,2 linkages in conjugated Le^b/y^ epitopes (Supplementary Fig. 6c,d), which concurred with our MS-based analyses on mucin.

In summary, *Am*GH29C and *Am*GH95B resulted in highest overall reduction of α1,3/4- as well as α1,2-fucosylation, respectively (Supplementary Table 4) and highest relative activities on HMOs and mucins (Supplementary Table 5). The broad specificity of *Am*GH29C is illustrated by activity on internal fucosylation and double sulphation, whereas *Am*GH95B was distinguished by activity on all H epitopes and double fucosylation. Collectively, the fucosidase suite allows full removal of mucin fucosyl substituents including from highly sulphated motifs.

### *A. muciniphila* sialidases remove all known mucin sialic acid linkages and include a sialyl-T-antigen-specific enzyme

The *Am*GH33A (Amuc_0625) and *Am*GH33B (Amuc_1835) are assigned into the CAZy family GH33. Two additional sequences (Amuc_0623 and Amuc_1547), with bacterial-neuraminidase-repeat-like domains that form the catalytic β-propeller fold in GH33 sialidases (Supplementary Fig. 7a), were previously reported as active sialidases, based on an indirect chromogenic assay^25^. Sequences of Amuc_1547 and its close orthologues populate a separate and distant cluster in the GH33 phylogenetic tree (Supplementary Fig. 7b,c), which together with substitutions in the catalytic residues (detailed below) justified the enzyme encoded by Amuc_1547 (*Am*GHxxx) to be the defining member of the new GHxxx family.

We expressed all four enzymes and assayed their activity on 3’-sialyl lactose (3sL), 6’-sialyl lactose (6sL), sialyl-Le^a^ and on α2,8-polysialic acid oligomers, which revealed activity for *Am*GH33A, *Am*GH33B and *Am*GHxxx, but not Amuc_0623 (Supplementary Fig. 8).

Next, we evaluated the four *A. muciniphila* sialidases against released *N*-glycans from human immunoglobulin G, released *O*-glycans and from PCM, and intact Muc2 from mouse (Muc2_Mouse_). Sialidase activity was observed for *Am*GH33A, *Am*GH33B and *Am*GHxxx on four abundant sialyl-motifs, but Amuc_0623 activity was not reliably measured (Fig. 2a-c). The active enzymes released both Neu5Ac and the animal-derived *N*-glycolylneuraminic acid (Neu5Gc), from porcine mucin (Fig. 2d-e). The motifs targeted by each enzyme were not dependent on the specific *O*-glycan core (Supplementary Fig. 9a-d).

**Fig. 2:**
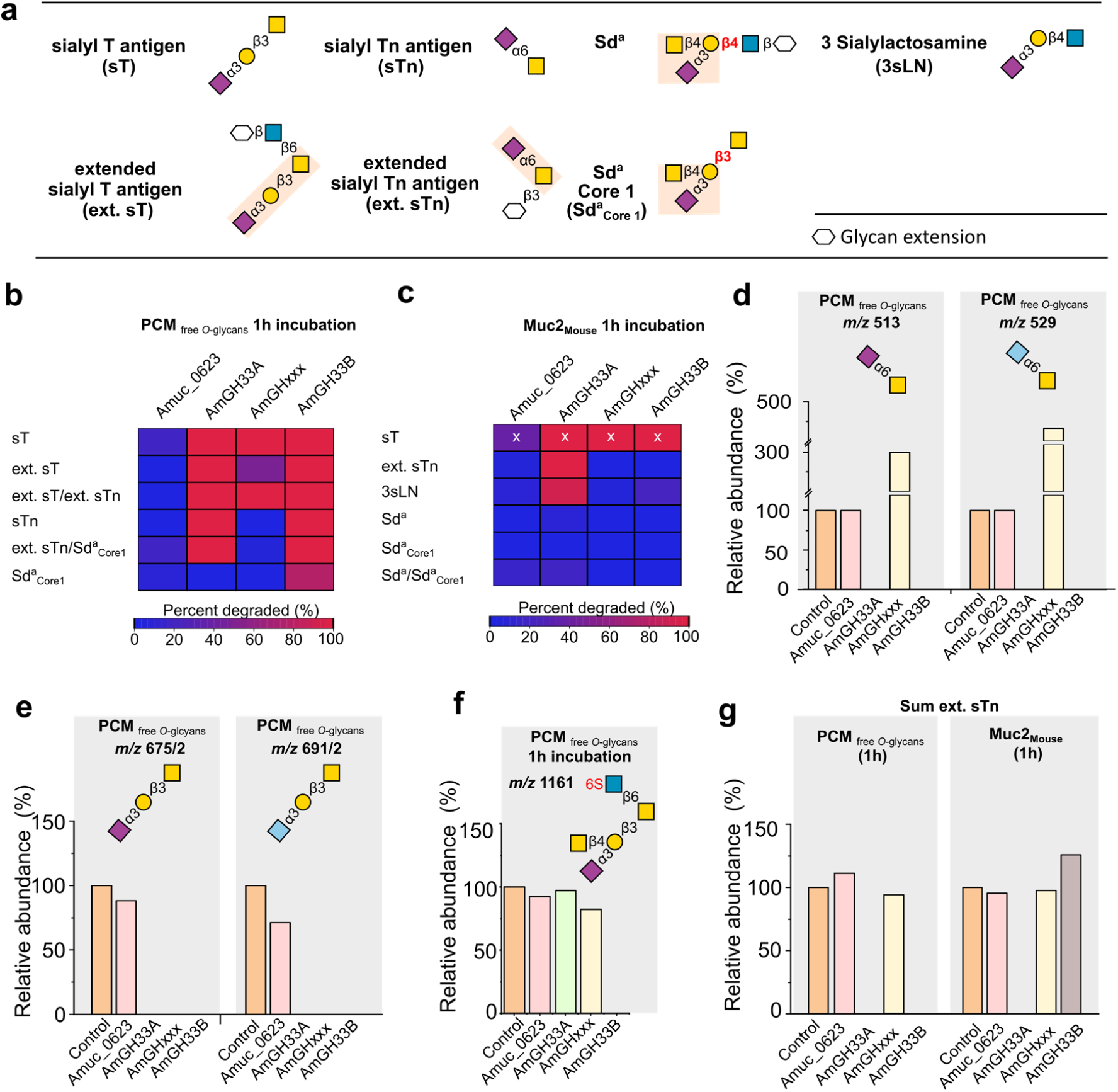
The activity profiles of *A. muciniphila* sialidases. **a**, Overview of sialylated epitopes, as denoted with a lower case “s”, in the analyzed mucins and released mucin *O*-glycans. **b**, and **c**, Activity heat maps on sialylated epitopes in released porcine colonic mucin (PCM) *O*-glycans and Muc2 from mouse (Muc2_Mouse_), respectively after a 1 h incubation. **d**, and **e**, Activity profiles on selected glycans revealing the broad substrate recognition of *Am*GH33A and *Am*GH33B towards α2,3- and α2,6-linked the Neu5Ac and Neu5Gc (light blue) sialic acid forms. **f**, Selected examples illustrating differences between *Am*GH33A and *Am*GH33B. **g**, A comparison of enzyme activity profiles on the sum of sialylated extended Tn epitopes either in free *O*-glycans from porcine colonic mucin (PCM) or attached *O*-glycans in Muc2_Mouse_. All data are from a single experiment. The “x” marked data are obtained from a single glycan structure, due to the low abundance of the sialyl T epitope in the Muc2_Mouse_ sample. Sulphatyl substitutions are denoted with capital S in red and their linkage is left out, if the assignment was not possible based on the MS data.

Both GH33 enzymes were active on sialyl decorations, *e*.*g*. in mucin α2,3- and α2,6-sialyl linkages on both *O*- and *N*-glycans (Fig. 2b-e; Supplementary Fig. 9e). Striking specificity differences, however, between the two enzymes were observed. Thus, *Am*GH33B was not hindered by the substitution of the galactosyl moiety of Neu5Acα2,3Gal, *e*.*g*. on Sd^a^_Core1_, as opposed to *Am*GH33A which was inactive on the same motif in released PCM *O*-glycans after 1 h incubation (Fig. 2a,b,f). By contrast, only *Am*GH33A displayed similar efficiency towards the ext. sTn epitope on both PCM released *O*-glycans and Muc2_Mouse_ attached *O*-glycans after 1 h reactions (Fig. 2a-c,g). Similarly, only *Am*GH33A cleaved Neu5Acα2,3Gal in both released PCM *O*-glycans and intact Muc2_Mouse_ after 1 h (Fig. 2b,c, Supplementary Fig. 9f). Our findings illustrate the importance of the sialyl density and glycan context (free/attached) in interrogating enzyme efficiencies on specific *O*-glycan motifs (Supplementary Tables 6 and 7), and merits caution when inferring enzyme specificities based on a single time point on free *O*-glycans.

Uniquely, *Am*GHxxx displayed exclusive specificity towards the sialyl-T-antigen, but accepts α2,6-sialyation or β1,6-substitution of the GalNAc unit (Fig. 2a-c,e,g; Supplementary Fig. 9g). Thus, *Am*GHxxx is inactive on a substituted Gal unit of the T-antigen, *e*.*g*. Sd^a^ epitopes (Fig. 2b,c,f), or on a different linkage/monosaccharides to the reducing end of the Neu5Acα2,3Gal motif, *e*.*g*. in 3sLN (Fig. 2b,c, Supplementary Fig. 9f). To our knowledge, this strict specificity is unprecedented amongst reported sialidases.

### *Am*GH29D adopts a “Cobra strike pose” architecture, previously not observed in fucosidases

Both of the *Am*GH29C/D enzymes consist of a catalytic N-terminal domain, followed by a predicted galactose binding like domain (GBLD), an unassigned sequence patch, and a C-terminal CBM32 (Supplementary Fig. 1a, Supplementary Fig. 2a,b). Amongst biochemically characterized enzymes, this architecture was only observed in the mucolytic specialists *Bifidobacterium bifidum*. Although crystallization attempts of both *Am*GH29C/D were carried out, we could only determine the structure of *Am*GH29D, the most complex fucosidase structure to date (Fig 3 a,b). Unique to this structure is that the GBLD domain is joined to a linker domain and a C-terminal CBM32 (Fig 3 a,b, Supplementary Fig. 10a). The linker domain and the CBM32 form an extended structure, which positions the CBM32 binding site above the catalytic domain (Fig 3 a,b). This juxtapositioning of the linker-CBM32 domain relative to the catalytic domain, which resembles a Cobra strike pose, is observed in the sialidase from *Micromonospora viridifaciens* (Supplementary Fig. 10a-c), suggesting a case of convergent evolution to a similar putative mucin-binding motif. The GBLD and the CBM32 share the same fold (Supplementary Fig. 10d), despite less than 22% shared sequence identity (Supplementary Fig. 1b). The position and the chemistry of the putative binding sites of the GBLD and the CBM32 domains are different, suggesting their possible functional divergence (Supplementary Fig. 10e-g). The catalytic site of *Am*GH29D is similar to the closest characterised counterpart from *Streptococcus pneumonia* (Supplementary Table 8), but differs by being flanked with a flat positively charged surface (Supplementary Fig. 11a), compatible with binding sialylated or sulphated glycans at the mucin surface.

**Fig. 3:**
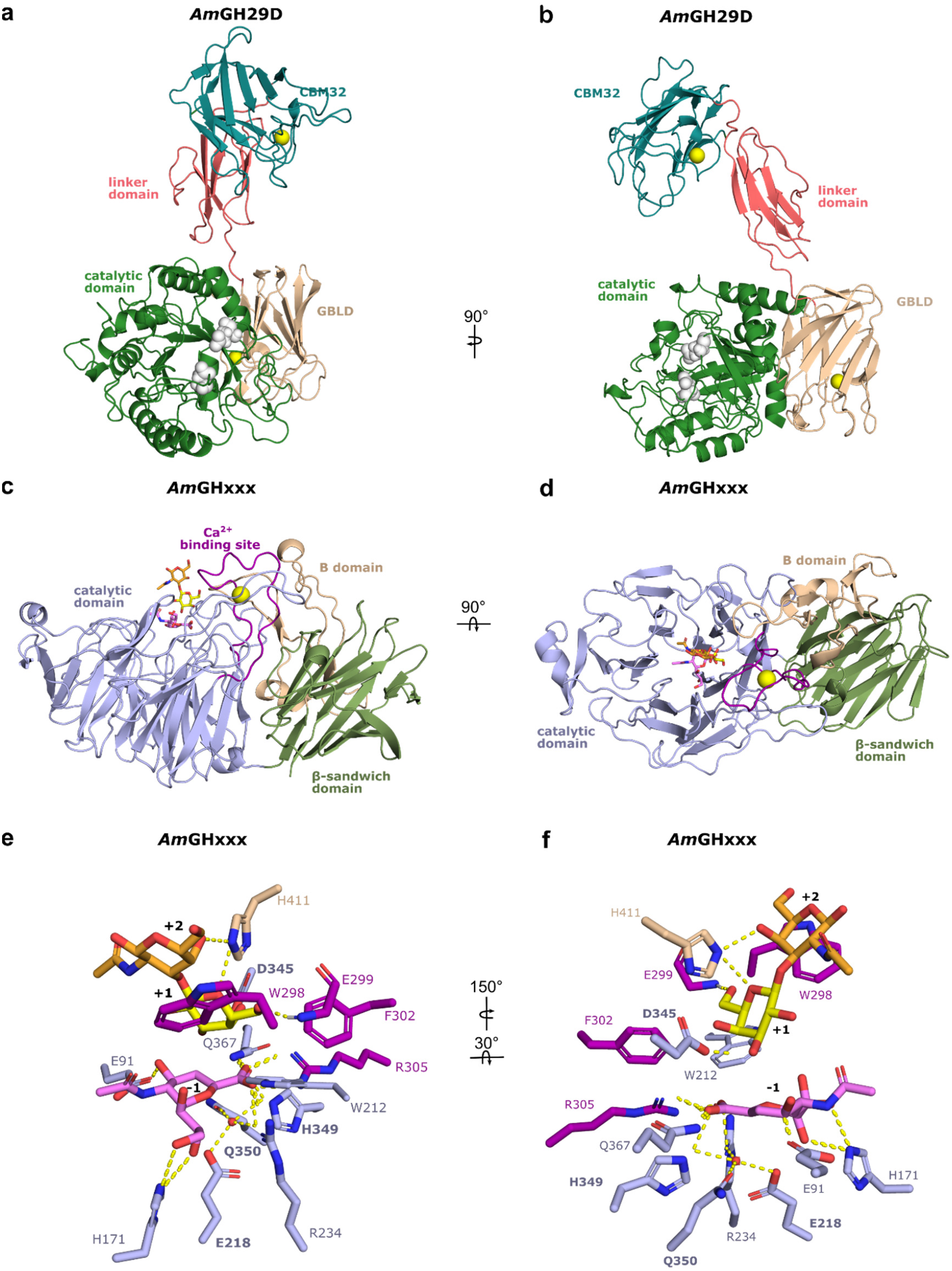
The crystal structures of the *Am*GH29D fucosidase and *Am*GHxxx sialidase from *A. muciniphila*. **a**, Overall structure of *Am*GH29D comprising a catalytic N-terminal (β/α)8 domain (amino acids 38-362), a predicted β-sandwich galactose binding like domain (GBLD, aa 363-489), a linker domain comprising two β-sheets formed by five antiparallel strands (aa 490-571) and a C-terminal CBM32 (aa 572-704). The inferred catalytic residues (white) and the bound Ca^2+^ ions (orange) are shown as spheres. **b**, A 90° rotation of the view in a. **c**, Overall structure of *Am*GHxxx, comprising an N-terminal 6-fold β-propeller catalytic domain (aa 23-281 and 307-384) with a Ca^2+^-binding domain, formed by an extended loop between the two inner β-strands in the propeller blade 2 (aa 282-306). The Ca^2+^ (orange sphere) is assigned based on coordination geometry and distance. The catalytic domain is joint to a C-terminal β-sandwich CBM-like domain (residues 457-595) and an inserted B domain (aa 391-456) between the β-strands 1 (sheet I) and 2 (sheet II) of the CBM-like domain. **d**, A 90° rotation of the view in c. **e**, The active site of *Am*GHxxx with the inhibitor *N*-Acetyl-2,3-dehydro-2-deoxyneuraminic acid (DANA) at subsite -1, and the Gal and GalNAc units of the T-antigen disaccharide bound at the +1 and +2 subsites, respectively. The same domain colours are used in panels c-f. The bold font highlights invariant potential catalytic residues in GHxxx.

To explain the specificity differences between *Am*GH29C and *Am*GH29D, we generated an AlphaFold model of *Am*GH29C based on *Am*GH29D. Shortening of loops as compared to *Am*GH29D (Supplementary Fig. 11d-e), results in a more open active site of *Am*GH29C. This allows the accommodation of sulphated or larger fucosylated substrates that are extended at non-reducing ends, consistent with exclusive activity of *Am*GH29C on internal fucosylated GlcNAc (Fig 1a-c,f; Supplementary Fig. 4k,n).

### *Structural signatures of the inverting mechanism and strict specificity of* Am*GHxxx*

To explain the strict specificity, we determined three structures of *Am*GHxxx, one in free form, one bound with the transition-state inhibitor DANA, and one bound to both the T-antigen disaccharide (GNB) and DANA, which provides a mimic for a substrate Michaelis complex. The structure comprises an N-terminal catalytic domain joint to a C-terminal CBM-like domain (Fig. 3 c,d). An inserted B domain between β-strands 1 (sheet I) and 2 (sheet II) of the CBM-like domain acts as a bridge by packing onto the catalytic domain via an extensive interface. A long loop in propeller blade 2 forms a Ca^2+^-binding domain (Fig. 3 c,d). The CBM-like domain occurs uniquely within GHxxx (Supplementary Fig. 12a), with only very distant structural similarities to Galectin galactoside-binding domains (Supplementary Table 10).

The catalytic module of *Am*GHxxx is distantly related to counterparts in GH33 (Supplementary Table 9). The catalytic site comprises a shallow pocket, flanked by a positive electrostatic potential (Supplementary Fig. 13a). The active site is open at one side of the β-propeller due to shorter loops as compared to GH33 enzymes. The Ca^2+^-binding domain, the B domain and two large loops pack onto the β-propeller to give the enzyme a “sun-chair” architecture (Fig. 3c,d; Supplementary Fig. 13b-f).

At the catalytic site, R234 and R305 are shared with GH33, whereas a glutamine (Q367) substitutes the third arginine in the GH33 conserved triad (Fig. 3e, f; Supplementary Fig. 14a). A unique signature is the substitution of the tyrosine catalytic nucleophile in the GH33 family to a glutamine (Q350) that is preceded by a histidine (H349) in *Am*GHxxx (Supplementary Fig. 14b-d). Strikingly, Q350 and an adjacent glutamate (E218), both invariant in GHxxx (Supplementary Fig. 12b c), are potentially hydrogen bonded to a water molecule that overlays with the oxygen in the GH33 catalytic tyrosine. This water is positioned for nucleophilic attack at the C2 of the sialyl (or the inhibitor) at subsite -1 (Fig 1e,f; Supplementary Fig. 14b). Notably, a solvent tunnel potentially provides catalytic water to the enzyme catalytic site (Supplementary Fig. 14e,f), similarly to some exoglycosidases^26^. An invariant aspartate (D345), unique for GHxxx, is hydrogen bonded to the Gal C3-OH group at subsite +1 (Supplementary Fig. 14b). Based on the data from the founding *Am*GHxxx, we propose, that the inverting mechanism of GHxxx, involves the activation of a water molecule to act as a nucleophile instead of a tyrosine in GH33. The conservation and the position of D324 (Supplementary Fig. 12b c) qualify this residue to be the catalytic acid in the mechanism.

To investigate the stereochemical mechanism, the hydrolysis time course of 3’-sialyllactose by *Am*GHxxx was analyzed using real-time NMR spectroscopy. The initially emerging signals were for β-Neu5Ac and α-Neu5Ac, in the reactions catalyzed by *Am*GHxxx and *Am*GH33A, respectively (Supplementary Fig. 15). These data corroborate the proposed inverting mechanism of *Am*GHxxx, contrasting the retaining GH33 family and the larger evolutionary related CAZy clan E.

Comparison of free and the ligand bound structures of *Am*GHxxx revealed the flipping of a tryptophan (W298) in the ligand-bound structures (Supplementary Fig. 16a,b). This conformational change positions W277 (56.3 % conserved within GHxxx, Supplementary Fig. 12c b), to stack onto the GalNAc unit of the T-antigen, thereby defining subsite +2. The invariant histidine (H411) from domain B forms potential hydrogen bonds to the Gal and GalNAc units. Thus, W298 and H390 form a “sugar tang” that restricts the T-antigen disaccharide. The galactosyl at subsite +1 stacks onto an invariant tryptophan (W212) and is additionally recognized by two hydrogen bonds (Fig. 3e, f, Supplementary Fig. 12c). Collectively, these aromatic stacking and polar interactions provide a plausible explanation for the strict specificity of the enzyme.

The presence of two potential saccharide surface binding sites (SBSs) is an intriguing observation. The first of these sites is adjacent to the active site, where an inhibitor molecule was modelled, whereas a galactosyl unit was modelled at the second binding site located on the opposite side of the catalytic site (Supplementary Fig. 16c-g). Although both SBSs are conserved in *Akkermansia* sequences that populate a single clade in the phylogenetic tree of GHxxx, only moderate conservation was observed for other phylogenetic clusters (Supplementary Fig. 12d-g). The presence of a CBM-like domain and of two potential secondary binding sites are indicative of the association of the enzyme to mucin, which is explored below.

### Key fucosidases and sialidases display mucin binding and their corresponding activities are cell attached

The presence of putative CBMs prompted us to investigate enzyme binding to PGM. Strikingly, *Am*GH29C, *Am*GH29D, *Am*GH33B and *Am*GHxxx, all containing annotated or putative CBMs, were mainly bound to mucin in pull-down assays (Supplementary Fig. 17a,c-d), whereas no binding or weak binding (*e*.*g*. GH95s, Supplementary Fig. 17b,d) was shown for the rest of the enzymes. To test localization, we grew cells on PCM and assayed the intact cells and cell lysates against 2’FL (GH95 substrate), Le^a^ trisaccharide (*Am*GH29C/*Am*GH29D substrate), the disaccharide Fucα1,3GlcNAc (*Am*GH29A/*Am*GH29B substrate) and 6sL (*Am*GH33A/*Am*GH33B substrate). Enzymatic activity on 2’FL, Le^a^ and 6sL, but not Fucα1,3GlcNAc, was detected mainly in the cell fraction (Supplementary Fig. 18), whereas no activity was detected in the cell lysate. Enzymatic activity was also detected in the supernatant, especially at higher OD values and longer incubation times, which could be due to enzyme release due to proteolysis. These results are supportive of the display of the corresponding sialidases and fucosidases on the surface of *A. muciniphila* and not in the periplasm.

### *A. muciniphila* fucosidases and sialidases are crucial for growth on mucin and sharing of mucin derived sugars

In the absence of gene knock-out tools for *A. muciniphila*, we deployed inhibitors to probe the impact of the fucosidases and sialidases on mucin *O*-glycan utilization. First, we evaluated the inhibition potency toward the fucosidases (*IC*_50_<54 μM) and sialidases (*IC*_50_<200 μM) (Supplementary Tables 11 and 12). *A. muciniphila* cells grew undistinguishably on monosaccharides in presence or absence of 1 mM (Fig. 4a) or 20 mM (Fig. 4b) of each inhibitor. By contrast, growth on mucin (PCM) in the presence of the inhibitors was heavily impaired at 1 mM and completely abolished at 20 mM of each inhibitor (Fig. 4a,b; Supplementary Table 13). The results showed that the fucosidase and sialidase suite was critical for initiating mucin deconstruction. Next, we compared the growth of *A. muciniphila* on the intact and de-sialylated/de-fucosylated PCM. Strikingly, the difference in growth profiles was modest (Fig. 4c), indicating that the rate of release of fucose/sialic acid was not the limiting step during growth on mucin. Growth assays showed that sialic acid did not sustain *A. muciniphila* growth, whereas fucose supported very slow growth, in agreement with previous studies^27, 28^ (Fig. 4d). The lack of relevant contribution of the released sialic acid and fucose during growth on mucin highlighted possible nutritional sharing of these monosaccharides with other GM. We performed co-cultures of *A. muciniphila* and other mucus-associated butyrate-producing Clostridia to investigate cross-feeding. Generally, the tested strains grew poorly on mucin, consistent with low butyrate levels in culture supernatants (Fig. 4d-n). Significantly higher butyrate concentrations, however, were observed in co-culture supernatants of all but one strain with *A. muciniphila* (Fig. 4e,g, i and m), which was suggestive of nutritional sharing by *A. muciniphila* to mucus-associated Clostridia. Notably, the butyrate production was highest for *Roseburia inulinivorans* and *Faecalibacterium prausnitzii*, in excellent agreement with the growth of both species on sialic acid (Supplementary Fig. 19c). These findings suggest that the decapping enzymes promote direct sharing of their product monosaccharides to other GM, besides their key role for initiating mucin deconstruction and the access of *A. muciniphila* to the underlying glycans.

**Fig. 4:**
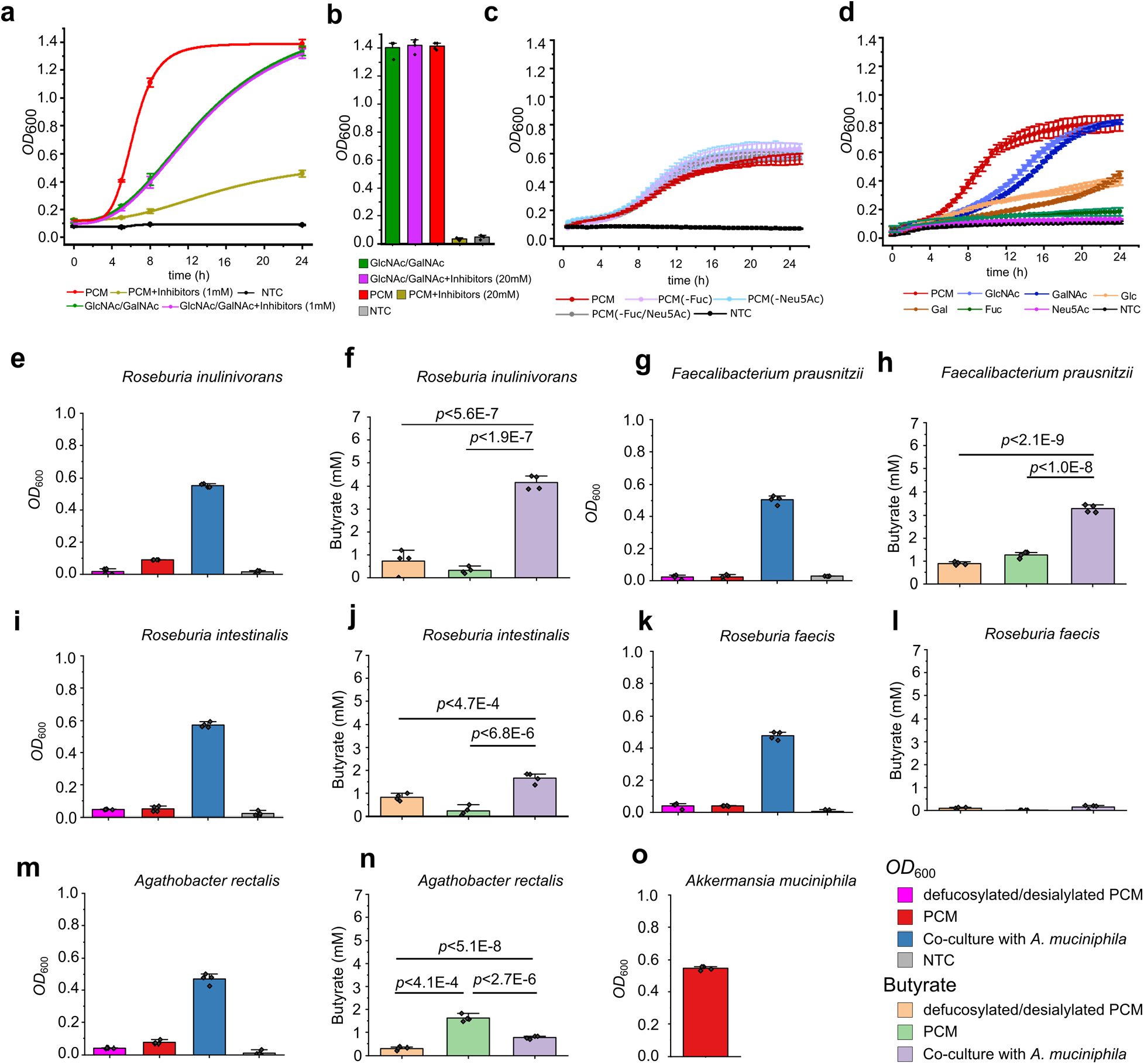
Fucosidases and sialidases of *A. muciniphila* are critical for growth on mucin and contribute to nutrient sharing with butyrate producing Clostridia. **a**, Growth curves of *A. muciniphila* on porcine colonic mucin (PCM) and on an equimolar mixture of GlcNAc/GalNAc, alone or in the presence of a equimolar mixture of fucosidase (1-Deoxyfuconojirimycin HCl, DFJ) and sialidase (*N*-Acetyl-2,3-dehydro-2-deoxyneuraminic acid, DANA) inhibitors (1mM each) as compared to no-carbon source control (NTC). **b**, Same as in a, but *A. muciniphila* growth is showed after 24 in the presence or absence of 20 mM of each inhibitor. **c**, Growth curves of *A. muciniphila* on PCM, de-fucosylated PCM and de-sialylated PCM compared to a no-carbon source control. The growth experiment is performed in a microtiter plate as opposed to panel a, where the growth was performed in a larger volume and the *OD*_600_ was measured in cuvettes. **d**, Growth curves of *A. muciniphila* on monosaccharides from mucin and a no-carbon source control. **e, g, i**,**k, and m**, Growth levels of different butyrate producing Clostridia from the Oscillospiraceae and Lachnospiraceae families on PCM and de-fucosylated/de-sialylated PCM (including a no-carbon source control) in monocultures or in co-culture with *A. muciniphila* on PCM after 24h. **f, h, j**, and **n**, The corresponding butyrate concentrations in culture supernatants. **o**, Growth level of *A. muciniphila* in monoculture on PCM after 24h. Growth analyses (a-c, d, f, h, j, l and n) on media supplemented with 0.5% (w/v) mucin or a 1:1 mix of GlcNAc and GalNAc, were performed in four independent biological replicates (n=4) and butyrate in culture supernatants was analyzed from 3 independent growth experiments (n=3). The growth data and butyrate quantifications are mean values with standard deviations (SD) as error bars.

## Discussion

The modulation of the metabolic and immune systems of the host by *A. muciniphila*^29, 30^ correlates to the relative abundance of this symbiont and its interplay with other mucus-associated bacteria. Remarkably, the molecular mechanisms of mucin glycan degradation^31^ and utilization by *A. muciniphila* remain largely underexplored. Here, we present detailed enzymatic, microbiological and structural analyses, which led to the identification of key glycoside hydrolases that collectively were able to removal all known mucin fucose and sialic acid caps, from mucin *O*-glycans.

The prevalence of the genes encoding the key fucosidases and sialidases in different *A. muciniphila* strains (Supplementary Table 14), underscores the importance of these enzymes for mucin *O*-glycan utilization. In addition, the sequence divergence of catalytic modules is consistent with the observed distinct enzymatic signature of each enzyme.

Interestingly, two of the enzymes, *Am*GH95A and *Am*GHxxx, display exclusive specificities towards a single glycan each, *e*.*g*. the abundant H2^32^ and the sialyl-T-antigen epitopes, respectively. To our knowledge, *Am*GH95A and *Am*GHxxx, are the most specific fucosidase and sialidase reported to date. By comparison, *Am*GH95B and *Am*GH29C are highly promiscuous, enabling the removal of multiple complex epitopes, *e*.*g. Am*GH95B targets H1, H2, H3 type and Le^b/y^ antigens (Fig. 1b, i-k) and *Am*GH29C is active on internal fucosylations, sialylated Le^a/x^ and Le^b/y^ epitopes as well as double sulphated motifs (Fig. 1b-h, Supplementary Table 4). The previously not reported ability to hydrolyze fucose from double sulphated motifs, suggests that de-fucosylation by *A. muciniphila* is feasible without prior de-sulphation.

The evolution of a specificity gradient that spans mono-epitope-specific to highly promiscuous fucosidases and sialidases, may be driven by the optimization of enzyme affinities (*K*_m_) to promote efficient decapping of highly diverse mucin *O*-glycans to promote growth. The importance of enzyme affinity is also evident from the strong association of *Am*GH29C, *Am*GH29D, *Am*GH33B and *Am*GHxxx to mucin. Mucin-binding of these enzymes correlates to the presence of putative carbohydrate-binding modules and saccharide surface binding sites, both known to increase the enzymatic efficiency of CAZymes towards insoluble glycans^33, 34^, which also applies to mucus matrix.

The observed extracellular cell-attached localization of fucosidase and sialidase activities indicate that sialic acid and fucose removal occurs fully extracellularly, which is crucial for further mucin glycan breakdown by the extracellular exo-glycosidases and endo-glycanases of *A. muciniphila* with ^31^. This strategy is in excellent agreement with impairment of *A. muciniphila* growth on PCM in the presence of sialidase and fucosidase inhibitors (Fig. 4a,b) and with a critical role of the decapping enzymes in initiating mucin deconstruction. *Bifidobacterium bifidum* extracellular fucosidases and sialidases mediate cross-feeding on mucin with other infant gut bifidobacteria^35^. Here we show that *A. muciniphila* may assume a similar ecological role amongst the mucus-adherent microbial community in adults, which is dominated by Lachnospiraceae, Oscillospiraceae (Fig. 4d-m) and to a less extent *Bacteroides* species^36, 37^. The butyrate concentration was commensurate to the growth of co-cultured Clostridia on sialic acid (Supplementary Fig. 19), which unveils the important contribution of this monosaccharide to the altruistic behavior of *A. muciniphila*^38, 39^. The very poor growth of *A. muciniphila* on fucose (Fig. 4d) is also expected to allow sharing this monosaccharide with fucose-utilizing bacteria^40^.

*Bacteroide*s species typically rely on minimal extracellular processing^12^ and subsequent import of large complex glycans for further periplasmic deconstruction^41^. By contrast, extracellular mucin decapping by *A. muciniphila*, is compatible with leakage of released sugars to mucus-associated Clostridia that possess efficient ABC transporters of fucosylated-glycans and sialic acid^39, 42^. The “soft” unselfish strategy is consistent with the observed cross-feeding (Fig. 4e-n) and may partially explain the poor competitiveness of *A. muciniphila* upon increased abundance of other mucolytic Clostridia, *e*.*g. Ruminococcus* species during dysbiosis^17^. Our work brings unique insight into the initial steps of mucin *O*-glycan deconstruction by *A. muciniphila* and on the mechanism of nutrient sharing with major mucus-associated butyrogenic Clostridia.

### Conclusions

Microbial mucin turnover is of paramount significance for the maintenance of symbiosis between the host and the mucus-associated microbiota as well as for pathologies linked to the excessive breakdown of the mucosal layer. Our findings offer unprecedented insight into the enzymatic apparatus that initiates mucin degradation and promotes nutrients sharing by a key dedicated mucolytic symbiont with the mucus-associated microbiota. The identification of the key enzymatic activities responsible for de-shielding mucin offers possible targets to block mucin collapse to treat downstream inflammatory disorders. The exquisite specificity of *A. muciniphila* sialidases and fucosidases expands the analytical toolbox for unambiguous linkage assignment in MS-based *O*-glycan analyses or for targeting specific glycan motifs.

## Materials and Methods

### Chemicals and carbohydrates

*p*-Nitrophenyl α-L-fucopyranoside (*p*NPFuc), *N*-acetylneuraminic acid (Neu5Ac), α-L-fucose (Fuc), Galactose (Gal), Glucose (Glc), *N*-acetylglucosamine (GlcNAc), *N*-acetylgalactosamine (GalNAc), Fetuin (from fetal bovine serum) and Type III porcine gastric mucin were from Sigma (St. Louis, MI, USA). Lewis antigens (Le^a^ triose, Le^x^ triose, Le^b^ tetraose, Le^y^ tetraose), 6-Sialyllactose (6sL), 3-Sialyllactose (3sL), 2’-Fucosyllactose (2FL), 3-Fucosyllactose (3FL) were purchased from Dextra (Reading, UK). αFuc1,3GalNAc, αFuc1,4GalNAc, αFuc1,2Gal and αFuc1,3Gal were from Toronto Research Chemicals (Toronto, Canada), Lacto-*N*-difucopentaose I (LNDFP I), Lacto-*N*-difucopentaose II (LNDFP II), Lacto-*N*-difucohexaose I (LNDFH I), Lacto-*N*-difucohexaose II (LNDFH II), sialylated Le^a^ triose (sLe^a^), Colominic acid, *N*-Acetyl-2,3-dehydro-2-deoxyneuraminic acid sodium salt (DANA), 1-Deoxyfuconojirimycin HCl (DFJ), 4-Methylumbelliferyl *N*-acetyl-α-D-neuraminic acid sodium salt (4MU-Neu5Ac) and Galacto-*N*-biose (GNB) were from Carbosynth (Berkshire, UK). Recombinant P-selectin glycoprotein ligand-1 (PSGL-1) with *O*-glycans that are terminated with Lewis antigens were prepared in Gothenburg University. Antibodies against Lewis antigens were from Sigma and Santa Cruz biotechnology. All purchased chemicals were of analytical grade unless otherwise stated and were used without further purification.

### Porcine gastric and colonic mucins and mouse colonic mucin

Type III porcine gastric mucin (PGM), purchased from Sigma (St. Louise, MI, USA), was further purified according to Miller et al^43^. Briefly, 2.5% (w/v) mucin was dissolved in phosphate buffered saline (PBS) pH=7.2 and stirred (20 h, room temperature), followed by centrifugation (10,000x *g*, 30 min, 4°C. The soluble mucin-containing supernatant was collected and precipitated by adding ice-cold EtOH to 60% (v/v) twice. Thereafter, the purified soluble mucin was dialyzed against Milli-Q using a 50 kDa molecular weight cut off membrane (SpectraPore7, Rancho Dominguez, CA, USA), freeze dried, and subsequently stored at -20°C until further usex Porcine colonic mucin (PCM) and Muc2 from mouse (Muc2_Mouse_) were prepared and purified as previously described^39, 44^.

### Preparation of *O*-glycan from PGM and PCM and *N*-glycans from human IgG

*O*-glycans from PCM and PGM were released from mucin glycoproteins by reductive *β*-elimination, before glycans were desalted and dried as previously described^45^. *N*-glycans were released from serum human IgG (Sigma-Aldrich) using PNGase F (CarboClip, Spain). In short, human IgG (1 mg) was reduced (10 mM DTT, 95 °C, 20 min) and alkylated (25 mM iodoacetamide, in dark at RT for 1 h). The *N*-glycans were then released by PNGase F in 50 mM NH_4_OAc (pH 8.4), 37 °C overnight incubation, before *N*-glycans were reduced (0.5 M NaBH_4_ in 20 mM NaOH, 50 °C overnight), desalted and dried^45^.

### Cloning, expression and purification of putative fucosidases and sialidases

The gene fragments of the glycoside hydrolase families GH29, GH95, GH33 and the putative sialidases with Bacterial-Neuraminidase-Repeat (BNR)-like domains, which encode the mature peptides lacking the signal peptides (as predicted by SignalP 5.0)^46^, were amplified from *Akkermansia muciniphila* ATCC BAA-835 (DSM 22959) genomic DNA using the primers as shown in (Supplementary Table 1). Infusion cloning (Clonetech/takara, CA, USA) was used to clone these amplicons into the NcoI and XhoI sites of the pET28a(+) vector (Novagen, Madison, WI). The resulting recombinant plasmids, which encode the members of GH29, GH33 and GH95 enzymes were transformed into *Escherichia coli (E. coli)* DH5α and transformants were selected on LB plates supplemented with kanamycin (50 µg mL^-1^). After full sequencing, the plasmids were transformed into a *E. coli* BL21 (DE3) Δ*lacZ* production strain (a kind gift from Professor Takane Katayama, Kyoto University, Kyoto, Japan) and the transformants were grown in 2 L LB medium with kanamycin (50 µg mL^-1^) at 30 °C to *OD*_600_≈0.5, followed by cooling on ice for 30 min before induction with isopropyl β-D-1-thiogalactopyranoside (IPTG) to 200 µM. Thereafter, growth was continued overnight at 18 °C and cells were harvested by centrifugation (10,000 *g*, 30 min), re-suspended in 10 ml of the purification buffer A (20mM HEPES, 500 mM NaCl, 10mM imidazole, 10% (v/v) glycerol, pH 7.5) and disintegrated by one passage through a high pressure homogenizer (SPCH-1, Stansted Fluid Power, Essex, UK) at 1000 bar, followed by incubation for 30 min on ice with 5 µl benzonase nuclease (Sigma). The lysates were then centrifuged for 20 min at 45,000 *g* and 4 °C and the supernatants were filtered (0.45 μM) and loaded onto HisTrap HP columns (GE Healthcare, Uppsala, Sweden). Then, bound proteins were washed (13 column volumes, CV) and eluted with the same buffer using an imidazole gradient from 10 to 400 mM in 15 CV. Pure protein fractions based on SDS-PAGE analysis were collected, concentrated, applied onto a HiLoad 16/600 Superdex 75 prep grade column (GE healthcare) and eluted by 1.2 column volumes of 20 mM HEPES, 150 mM NaCl, pH 6.8. The fractions containing each enzyme were pooled and concentrated (10 kDa Amicon® Ultra Centrifugal filters, Millipore, Darmstadt, Germany). Pure fractions, as judged by SDS-PAGE analysis, were pooled, concentrated as above and the protein concentration was determined using a Nanodrop (Thermo, Waltham, MA) using the theoretically predicted extinction coefficients at 280 nm (ε_280_) using the ProtParam tool (http://web.expasy.org/protparam). Finally, NaN_3_ (0.005% w/v) was added to the enzyme stocks that were stored at 4°C for further use.

### Enzyme activity and inhibition assays on synthetic substrates

All enzyme activity and inhibition reactions were performed in 20 mM HEPES, 150 mM NaCl, pH 6.8 unless otherwise state. For fucosidase kinetics, the initial reaction rates of *Am*GH29A (0.5 nM), *Am*GH29B (20 nM), *Am*GH29C (400 nM) *Am*GH29D (250 nM), *Am*GH95A (10 nM) and *Am*GH29B (50 nM) were determined on seven *p*NPFuc concentrations in the range 0.25-15 mM (except for *Am*GH95B, which was extended with a 30 mM substrate concentration). The reactions were carried out at 37 °C for 3 hours for all enzymes except *Am*GH29D, which was incubated for 4 hours. Aliquots of the reactions were collected at 30 min and 40 min intervals for 3 h and 4 h reactions, respectively, and quenched into Na_2_CO_3_ (0.4 M final concentration). The concentration of the *p*NP enzymatic product was determined by measuring *A*_405 nm_ using a 96-well plate reader (BMG Labtech, Ortenberg, Germany) using a *p*NP standard curve (0 to 140 µM *p*NP). The Michaelis-Menten equation was fit to the initial rates using Prism 6 (GraphPad San Diego, USA). For determining the inhibition constants (*IC*_50_), reactions were performed continuously at 37 °C for 30 min in a microtiter plate and absorbance (*A*_405 nm_, fucosidases) or emission (*E*_450 nm;_ *E*xcitation_370 nm_, sialidases) was measured in 60 sec intervals using a 96-well plate reader. The initial reactions rates of *Am*GH95A (0.25 µM), *Am*GH95B (0.5 µM), *Am*GH29A (0.5 µM) *Am*GH29B (0.5 µM) *Am*GH29C (10 µM) and *Am*GH29D (10 µM) were determined using 2 mM *p*NPFuc and Deoxyfuconojirimycin (DFJ) over a concentrations range of 0.1-10 mM (*Am*GH29A, *Am*GH29B and *Am*GH29C) or 1-100 mM (*Am*GH95A, *Am*GH95B and *Am*GH29D). Sialidase inhibition reactions were determined at enzyme concentration of 50 nM, expect Amuc_0623 which was assayed at 200 nM. The initial rates were determined using 1 mM 4-Methylumbelliferyl *N*-acetyl-α-D-neuraminic acid (4MU-Neu5Ac) and 0.01–1 mM *N*-Acetyl-2,3-dehydro-2-deoxyneuraminic acid (DANA). A Hill equation was fit to the initial rates using OriginPro 2021. All enzyme activity and inhibition reactions were performed in independent triplicates.

### Thin-layer chromatography

Thin layer chromatography (TLC) was used to screen the specificity of GH29 and GH95 enzymes towards the fuco-oligosaccharides α-Fuc(1,3)GalNAc, α-Fuc(1,4)GalNAc, α-Fuc(1,3)Gal, α-Fuc(1,2)Gal, 2’FL, 3FL, Le^a^ triose, Le^x^ triose, Le^b^ tetraose, LNDFH I, LNDFH II, LNFP II, LNFP V and sialylated Le^a^ triose (sLe^a^ triose) while the GH33, GHxxx and Amuc_0623 putative sialidase where screened on 6sL, 3sL and Colominic acid. Reactions (10 μL) were carried out using 2 mM of each substrate, 0.5 μM of each enzyme in 20 mM HEPES, 150 mM NaCl pH 6.8 at 37 °C for 1 h. Aliquots of 2μL were spotted on a silica gel 60 F254 plate (Merck, Germany) and the products were separated using a mobile phase of butanol/ethanol/Milli-Q (5:3:2, v/v/v) except for products obtained from 3sL that were separated using a mobile phase of isopropanol/ethyl acetate/Milli-Q (3:2:1, v/v/v). The plates were dried, sprayed with 2 % 5-methylresorcinol, 80 % EtOH and 10 % H_2_SO4, all v/v, and visualized by tarring at 300 °C. All enzyme activity reactions analyzed by thin-layer chromatography were performed in independent triplicates.

### Enzymatic analysis towards recombinant P-selectin glycoprotein ligand-1 (PSGL-1)/immunoglobulin mIgG2b chimeras carrying defined Lewis epitopes

To demonstrate enzymatic activity against intact mucin-type glycoproteins, PSGL-1/mIgG2b chimeras were produced and purified in glyco-engineered CHO cells as previously described^47^. In short, the extracellular portion of PSGL-1 was genetically fused with mouse immunoglobulin G2b creating the PSGL-1/mIgG2b expression plasmid, which was expressed in CHO cells together with plasmids encoding *O*-glycan core enzymes and combinations of fucosyl transferase genes. Thus, CHO cells were programmed to express the Lewis antigens (Le^a^, Le^x^, Le^b^ or Le^y^, Supplementary Fig. 6) on the mucin-type fusion protein. The produced PSGL-1/mIgG2b were purified form the cell culture supernatants using goat anti-mouse IgG agarose beads (Sigma–Aldrich). Each enzyme (2 µM) was incubated with beads carrying PSGL-1 glycoprotein (displaying a distinct Le antigens) in 20 mM HEPES buffer 150 mM NaCl pH 6.8 at 37 °C for 3 h in 50 µl. The beads were boiled in presence of SDS-loading buffer containing 25 mM DTT for 10 min at 95°C. The samples were electrophoretically separated on 8% NuPAGE gels (Invitrogen, Waltham, MA, USA) and thereafter blotted onto PVDF membranes (Immobilon P membranes, 0.45 μM) according to the manufactures manual (Invitrogen). The membrane blots were blocked with phosphate-buffered saline containing 0.2% Tween-20 (v/v, PBS-T) and 3% bovine serum albumin (w/v, BSA), which was also used for the dilution of the following primary mouse antibodies, followed by washing with PBS-T twice for 5 minutes. Then each membrane was incubated with the corresponding primary antibody (mouse IgG anti-Lewis antigen, Sigma–Aldrich, 1:500 dilution) for 1h at 4°C, washed as above and then HRP-conjugated poly-clonal goat anti-mouse IgM (1:5000 dilution, Sigma–Aldrich) was added for 1 h at 4 °C and lastly washed with PBS-T. Bound secondary antibodies detected by chemiluminescence using the ECL kit according to the manufacturer’s instructions (GE Healthcare, Uppsala, Sweden). Finally, membranes were stripped with Restore Western Blot Stripping Buffer (Thermo Scientific) under agitation at room temperature for 20 min and re-probed with HRP-conjugated poly-clonal goat anti-mouse IgG Fc (1:5000 dilution, Sigma–Aldrich) for checking the integrity of the mouse IgG2b Fc domain of the fusion protein, then the bound antibody was visualized as above. The following primary mouse antibodies were used: IgG anti-Blood Group Lewis A (7LE) (Santa Group Biotechnology, cat log: sc-51512), IgM anti-Blood Group Lewis B (T218) (Santa Group Biotechnology, cat log: sc-59470), IgM anti-Lewis X antibody: CD15 (C3D-1) (Santa Cruz Biotechnology, sc-19648), IgM anti-Blood Group Lewis Y (F3) (abcam, cat log: ab 3359). The secondary goat antibodies for Lewis A: Peroxidase goat anti-mouse IgG, F(ab’)2 (Jackson ImmunoResearch, cat log: 115-035-006) and for Lewis B, X and Y: Peroxidase goat anti-mouse IgM antibody (Sigma-Aldrich A-8786).

### Enzymatic assay towards glycans from mucin or glycoprotein using ESI-LC MS/MS

Fucosidase activity of GH29 and GH95 enzymes was analyzed on a mixture of intact PCM, PGM and fetuin dot-blotted on PVDF membranes or on previously release *N*-glycans (from human IgG) dot-blotted on membranes. Sialidase activity was assayed using released *O*-glycans from PCM or PGM, Muc2_Mouse_ glycoproteins, or released *N*-glycans (human IgG) were used.

For dot-blot assays, whole mouse colonic mucin^48^ glycoproteins or released *O*- or *N*-glycans from porcine colonic mucin or from human IgG were transferred to PVDF membrane (Immobilon P membranes, 0.45 µm) using dot blotting apparatus separately (0.1 mg per dot). Each enzyme (50 µL, 1.5 µM in 20 mM HEPES buffer with 150 mM NaCl, pH 6.8) was incubated with the substrate dots. For analyzing GH29 and GH95 fucosidases, 24 h incubations were performed, while sialidase activity was tested in 1h (*O*-glycans from PCM and PGM, Muc2_Mouse_ glycoprotein) and 24 h (*O*-glycans from PCM and PGM, Muc2_Mouse_ glycoprotein and *N*-glycans from human IgG) incubations. Afterwards, the residual *O*-linked glycans on the dot were released by reductive β-elimination after rinsing. The released *O*-glycans were desalted, dried as described elsewhere^45^. The resultant glycans were purified by passage through graphitized carbon particles (Thermo Scientific) packed on top of a C18 Zip-tip (Millipore). Samples were eluted with 65% (v/v) ACN in 0.5% trifluoroacetic acid (v/v), dried, and stored at −20 °C until further enzymatic analyses.

Released glycans were resuspended in 10 μL of Milli-Q water and analyzed by liquid chromatograph-electrospray ionization tandem mass spectrometry (LC-ESI/MS) using a 10 cm × 250 µm I.D. column, packed with 5 µm porous graphitized carbon particles (Hypercarb, Thermo-Hypersil, Runcorn, UK)). Glycans were eluted using a linear gradient 0–40% acetonitrile in 10 mM NH_4_HCO_3_ over 40 min at a flow rate 10 µl min^-1^. The eluted *O*-glycans were detected using an LTQ mass spectrometer (Thermo Scientific, San José, CA) in negative-ion mode with an electrospray voltage of 3.5 kV, capillary voltage of -33.0 V and capillary temperature of 300 °C. Air was used as a sheath gas. Full scan (m/z 380-2000, two microscan, maximum 100 ms, target value of 30,000) was performed, followed by data-dependent MS^2^ scans (two microscans, maximum 100 ms, target value of 10,000) with normalized collision energy of 35%, isolation window of 2.5 units, activation ρ=0.25 and activation time 30 ms). The threshold for MS^2^ was set to 300 counts. The data were processed using Xcalibur software (version 2.0.7, Thermo Scientific). Glycans were identified from their MS/MS spectra by manual annotation as previously described^49^. Raw data was uploaded on Glycopost Glycopost (https://glycopost.glycosmos.org/preview/1939229808630f42efa8e7f). The peak area (the area under the curve, AUC) of glycan structure was calculated using the Progenesis QI software (Nonlinear Dynamics, Waters Corp., Milford, MA, USA). The AUC of each structure was normalized to the total AUC and expressed as a percentage.

### NMR spectroscopy

Substrate solutions (2.5 mM) of 6’-sialyllactose (for *Am*GH33B) or of 3’-sialyllactose (for *Am*GHxxx) were prepared in 50 mM MES buffer, pH 6.8 in ^2^H_2_O. A 200 μL aliquot of the substrate solution was transferred into a 3 mm NMR tube and the sample was placed into an 800 MHz Bruker Avance III instrument equipped with a 5 mm TCI cryoprobe and thermally equilibrated to 310 K. The sample was tuned, matched, and shimmed in order to allow a rapid monitoring of the *Am*GHXXX-catalyzed conversion. The reaction was started by the addition of AmGH17X (1 μL, 10 μM) or *Am*GH33B (1 μL, 10 μM) into the NMR tube to a final concentration of 50 nM and mixing briefly before starting the analysis. A time series of one-dimensional ^1^H NMR spectra was acquired to follow the reaction in real time. The ^1^H NMR spectra sampled 16384 complex data points for an acquisition time of the free induction decay of 1.28 seconds. For each time point, 16 transients were summed up with an inter-scan relaxation delay of 2.0 seconds and using two dummy scans per time point, resulting in a time resolution of approximately one min. To validate the assignment of the α-Neu5Ac, the reaction was restarted and an ^1^H-^1^H TOCSY (2048 × 256 complex data points sampling 123 ms and 16 ms in the direct and indirect dimension, respectively) was acquired using a 10 kHz spin lock field during a mixing time of 80 ms. The ^1^H-^1^H TOCSY spectrum on the reaction mixture containing intermediates of 6’-sialyllactose or 3’-sialyllactose reaction were compared with a reference standard spectrum of Neu5NAc. The NMR data were considered unambiguous when acquired in single time-series experiments. Restarted assays using ^1^H-^1^H TOCSY confirmed the interpretation. All NMR spectra were acquired and processed with ample zero filling using Bruker Topspin 3.5 pl7 software and were subsequently analyzed with the same software.

### Crystallization

The crystallization of *Am*GHxxx (Amuc_1547) was performed by the sitting drop method using a mosquito robot (mosquito Xtal3, SPT labtech, Melbourn, United Kingdom) to mix 0.15 μL reservoir: 0.15 μL enzyme (30.5 mg mL^-1^) and the plates were thereafter incubated at 18°C. The first crystals appeared after two weeks in the Index screen from Hampton Research (Aliso Viejo, CA, USA), condition 82 (0.2 M MgCl_2_·(H_2_O)_6_, 0.1 M BIS-TRIS pH 5.5, 25% w/v Polyethylene glycol 3,350). After optimization, the best crystals were obtained under the same condition as above, but using a lower concentration (18% w/v) of Polyethylene glycol 3,350. To evaluate if *O*-glycans from PCM could facilitate the crystallization of the enzyme with ligand, the crystallization protocol and condition as described above was used to co-crystallize *Am*GHxxx with PCM. Thus, the enzyme (30.5 mg mL^-1^) and PCM (4% (w/v) prepared in Milli-Q) were mixed at room temperature at a 1:1 ratio (v/v) (resulting in a final PCM concentration of 2% w/v) before the enzyme-glycan solution was mixed with the reservoir as described above and plates were incubated at 18 °C. Enzyme crystals were flash-frozen in liquid nitrogen in nylon loops using 25% ethylene glycol as cryoprotectant. Of note, the crystals from the co-crystallization appeared only after hours were much larger rhombus-shaped as compared to the crystals in the lack of added glycans.

Similarly, *Am*GH29D (25 mg mL^-1^) was co crystallized with the same glycan mixture as above and mixed with the glycan mixture (1:1 v/v). The first crystals appeared in the Molecular Dimensions structure screen 2 (Holland, OH, USA) condition 28 (0.1M HEPES pH 7.5, 20% PEG 10,000) at 18 °C. After optimization, the best diffraction data were obtained by mixing *Am*GH29D (25 mg mL^-1^) with glycans at 1:0.8 ratio (v/v) at room temperature, and reservoir condition 0.1M HEPES pH 7.7 16% PEG 10,000, and thereafter incubation of the plate at 16 °C. Diffraction data were collected at the BioMAX beamline at the MAX IV synchrotron radiation facility (Lund, Sweden) and the P13 EMBL Beamline at the DESY (Hamburg, Germany). The data was processed with XDS and the structures were refined using PHENIX.refine^50^ and manually rebuilt using Coot^51^ (Supplementary Tables 15 and 16). Structure validation was performed using MolProbity^52^.

### Fucosidase and sialidase activity measurements on whole cells and in culture supernatant

For localizing fucosidase and sialidase activity, *A. muciniphila* was grown in three biological triplicates anaerobically in 8 mL YCFA medium containing 0.5% (w/v) PCM for 16h. For preparing whole cells, 2 mL culture were harvested (5000 *g*, 10min at 4 °C) and cells were washed three times (5000 *g*, 10 min at 4 °C) with 1 mL ice cold 10 mM sodium phosphate, 150 mM NaCl, pH=6.5 buffer before resuspension to *OD*_600_=0.5 and *OD*_600_=8 in the same buffer. Cell lysates and cell debris were prepared by lysing (Qsonica sonicator, 5 mm probe tip, 4 × 15 s at 4 °C) *OD*_600_ adjusted cell suspensions (prepared as described above), separating insoluble cell debris from the clarified lysates (20,000 *g* for 30 min at 4 °C) and resuspension of the cell debris in the same buffer to equal volumes as the whole cell preparations to *OD*_600_=0.5 and *OD*_600_=8. For released proteins to culture supernatants, cells were removed (20,000 *g* for 20 min at 4 °C) from 2 mL cultures, the supernatants were exchanged three times to the same buffer as above (Amicon Ultra 0.5 mL, 10 kDa cut off (Merck, Darmstadt, Germany) (10,000 *g* for 20 min at 4 °C) before adjusting to the volumes of the whole cell preparation to *OD*_600_=0.5 and *OD*_600_=8. Next, thin layer chromatography was used to screen for fucosidase active towards 2FL, Le^a^ trisaccharide and Fuc(α1,3)GlcNAc as well as for sialidases activity towards 6sL. Reactions were initiated out by mixing 10 μL of 5 mM of each substrate in the same buffer as above and 10 μL whole cell, cell debris, cell lysate or supernatant solutions at 37 °C. Aliquots of 2μL were spotted on a silica gel 60 F254 plate (Merck, Germany) after 1 h, 2 h, 3 h, and 4 h and the products were separated, and the plates developed as described above. The growth assays were performed in three biological triplicates and a single TLC analysis was performed for each of the three biological triplicates.

### Mucin binding assay

Binding of *A. muciniphila* GH29/GH95 fucosidases and GH33/GHxxx sialidases to PGM and to Avicel (used as negative control) was assessed by a pull-down assay, followed by sodium dodecyl sulfate polyacrylamide gel electrophoresis (SDS-PAGE). In short, insoluble PGM and Avicel were washed three times (20,000 *g*, 5 min, 4 °C) with 1 mL standard buffer (20 mM HEPES buffer with 150 mM NaCl, pH 6.8), before resuspension to a concentration of 1 % (w/v) in the above buffer. Next, 50 μL of the PGM or Avicel suspensions were mixed with 50 μL of fucosidases (0.1 mg mL^-1^), sialidases (0.1 mg mL^-1^) or bovine serum albumin (0.1 mg mL^-1^) used as negative control), incubated for 20 min on 4 °C and centrifuged (20,000 *g*, 10 min, 4 °C). Resulting supernatants were transferred into fresh 1.5 mL reaction tubes and PGM/Avicel pellets were washed twice (20,000 *g*, 5 min, 4 °C) with 100 μL buffer, before resuspension in 100 μL standard buffer. Next, 100 μL protein solution was supplemented with 35 μL SDS sample buffer (NuPAGE) and the samples were boiled for 10 min, loaded (15 μL) into a gel, and analyzed using SDS-PAGE. The binding assay was performed in two independent replicates and SDS-PAGE analyses were performed once per independent replicate.

### Growth experiments, co-culture experiments and butyrate quantification

For single strain monocultures *Roseburia inulinivorans* DSM 16841, *Roseburia intestinalis* DSM 14610, *Roseburia faecis* DSM 16840, *Agathobacter rectalis* DSM 17629, *Faecalibacterium prausnitzii* DSM 17677 and *Akkermansia muciniphila* DSM 22959 were grown anaerobically at 37°C in YCFA media using a Whitley DG259 Anaerobic Workstation (Don Whitley Scientific). Growth media were supplemented with 0.5% (w/v) carbohydrates sterilized by filtration (soluble carbohydrates, 0.45 µm filters) or autoclaving (mucins, 15 min at 121 °C) and cultures were performed in at least three independent biological replicates unless otherwise indicated. For the inhibition of *A. muciniphila* fucosidases and sialidases, culture media were supplemented with the sterile filtered (0.45 µm filters) DFJ and DANA inhibitors to a final concentration of 1 mM or 20mM each. Bacterial growth was monitored by measuring *OD*_600_ and for growth experiments performed in airtight sealed microtiter plates (sealing tape for 96-well plates, Thermo Scientific), a Power Wave XS microplate reader (BioTek Agilent) was used to monitor *OD*_600_. For sialic acid and fucose quantification in culture supernatants aliquots (30 µL) were collected, mixed with 100 µL 0.9% (w/v) NaCl, before cells were removed by centrifugation (20,000 *g*, 4 °C 10 min). Next, supernatants were frozen at -20 °C before further analysis.

For co-culture experime*nts, R. inulinivorans, R. intestinalis, R. faecis, A. rectalis, F. prausnitzii* and *A. muciniphila* were grown in 10 mL YCFA to mid-late exponential phase (*OD*_600_=0.6-0.7). From these pre-cultures, equal amounts of cells (*OD*_600_) were used to inoculate 1 mL fresh YCFA medium supplemented with 0.5% (w/v) of PCM to a start *OD*_600_ =0.01. All cultures were performed in four independent biological replicates and growth was followed (*OD*_600_) by sampling at 0 and 24h. Samples (200 µL) from time 0 h and after 24 h were collected for SCFA quantification. The samples were centrifuged (20,000 *g*, 4 °C 10min) and the resulting supernatants diluted with 200 µL 5 mM H_2_SO_4_ before sterile filtrated (0.45 µm filters) and storage at −80 °C for further analysis.

Butyrate in culture supernatants was quantified as previously described^39^. In short, an HPLC coupled to a refracting index detector (RID) and diode array detector (DAD) on an Agilent HP 1100 system (Agilent) was used to quantify standards of butyric acid (0.09-25 mM) (prepared in 5 mM H_2_SO_4_) and 20 µL of standard or filtered (0.45 µM filter) culture supernatants samples from three biological replicates were injected on a 7.8 × 300 mm Aminex HPX-87H column (Biorad) combined with a 4.6 × 30 mm Cation H guard column (Biorad). Elution was performed with a constant flow rate of 0.6 mL min^−1^ and a mobile phase of 5 mM H_2_SO_4_. Standards were analyzed as above in three technical triplicates.

### Quantification and statistical analysis

For determining the statistical significance between butyrate concentrations in the different cultures, a one-Way ANOVA and Tukey post hoc test was used (OriginPro 2021). Statistical parameters, including values of n and *p*-values are reported in the figures and figure legends. The data are expressed as arithmetic means with standard deviations (SD), unless otherwise indicated. The statistical significance between growth levels reached on mucin/monosaccharides with and without the inhibitors was evaluated using an unpaired two-tailed Student’s *t*-test using OriginPro 2021.

### Bioinformatics

SignalP (v.5.0), PSORTb (v.3.0.3), TMHMM (v.2.0) were used for signal peptide and transmembrane domain prediction. CAZy, dbCAN meta server and InterPro were used under default settings to analyses size and modular organization of proteins. For phylogenetic analyses, sequences data sets were generated by identified orthologues via batch BLASTP searches using both *A. muciniphila* GH29, GH95 fucosidases or GH33/BRN repeat-like domain sialidases and those sequences that are defined as characterized in the particular GH families within the CAZy database as queries, against 7950 (meta)genomes from the human gut microbiota (retrieved from the PATRIC (v.3.6.12) database, date: November 2021, inclusion criteria: host: “human/*Homo sapiens*”, isolation source: “feces/fecal sample”, genome quality “good”). Redundancy in sequence datasets was reduced using CD-HIT server under default settings and with a sequence identity cut off = 0.95. Structural guided protein sequence alignments were performed using PROMALS3D and by using structurally characterized orthologues from the particular CAZy GH families. Phylogenetic trees were constructed using the MAFFT server interfaced (https://mafft.cbrc.jp/alignment/software/) (neighbor-joining algorithm and with bootstraps performed with 1000 replicates) and afterwards visualized in iTOL. The prevalence of the different enzymes was analysed using 177 *A. muciniphila* good quality genomes from the same database. Identification of closest structural characterized orthologues were done using the Dali server^53^. Orthologues to *Am*GHxxx were identified by a BlastP search against the nonredundant protein sequence and by using *Am*GHxxx (Amuc_1547) as query sequence. Redundancy of resulting sequencing (e-value cut off: 1e-25) was reduced the using CD-HIT server under default settings and with a sequence identity cut off = 0.90 and the redundancy reduced dataset was structurally guided aligned using the PROMALS3d server (used protein structures: *Am*GHxxx, and structurally characterized orthologues from the GH33). *Am*GHxxx orthologues were selected in resulting alignment by the presence of the conserved catalytic machinery as displayed by *Am*GHxxx. Alphafold modelling was performed using ColabFold^54^ on its web interface (https://colab.research.google.com/github/sokrypton/ColabFold/blob/main/AlphaFold2.ipynb) under standard settings (template mode on: (Structure of *Am*GH29D); msa_mode: MMseq2, pair_mode: unpaired and paired; model_type: auto, num_recycles: 3). Sequence logos were generated using the Seq2Logo^55^ web interface using standard settings (Logo type: Kullback-Leiber; Clustering method: Hobohm1; clustering threshold: 0.63 and 200 pseudo counts).

## Data availability

Data that support the findings of this study are available in the Article and the Supplementary material.

## Acknowledgements

The study was funded by the Ministry of Higher Education and Scientific Research of Iraq through a PhD scholarship for B.S. Additional funding was from the Independent Research Fund Denmark, Natural Sciences grant 1026-00386B for M.A.H. The NMR spectra were recorded at the NMR Center DTU, supported by the Villum Foundation. Saromics Biostructures AB (Dr. Maria Håkansson) are thanked for initial crystallization data. We acknowledge MAX IV Laboratory for time on Beamline BioMax under Proposal 20200120 Research conducted at MAX IV, a Swedish national user facility, is supported by the Swedish Research council under contract 2018-07152, the Swedish Governmental Agency for Innovation Systems under contract 2018-04969, and Formas under contract 2019-02496 and we acknowledge DESY (Hamburg, Germany), a member of the Helmholtz Association HGF, for the provision of experimental facilities. Parts of this research were carried out at P13 and we would like to thank Isabel Bento for assistance in data collection. Beam time was allocated for proposal(s) MX846.The authors would like to thank Professor Bernard Henrissat for his kind input and discussion on the defining member of GHxxx.

## Author contributions

B.S., E.N.K. and M.A.H. conceptualized the research and provided funding. M.A.H. led the study and wrote the manuscript with B.S. and M.J.P. B.S. cloned and produced the enzymes for the initial TLC and LC-MS analysis. B.S. and C.J. performed the enzymatic LC-MS analyses. B.S., M.J.P., C.J. and N.G.K. analyzed and interpreted the glycomic LC-MS data. B.S., J.L. and J.H. performed and interpreted the enzyme activity on recombinant PSGL1. H.W., A.M.G. and N.J. performed and analyzed the fucosidase kinetic assays. M.J.P. and S.M. designed the NMR analysis, which was carried out by S.M, who also generated the NMR analysis figure. M.J.P. performed the microbiology growth and inhibition assay, with help from T.S.N. The structural biology work was carried out by H.S., T.S.N. and J.P.M. The final version of the figures (except for the NMR analysis) was generated by M.J.P. All authors contributed to editing the manuscript and accept the present findings and conclusions.

