## Supplementary Material for "Sialidases and Fucosidases of *Akkermansia muciniphila* are crucial for growth on mucin and nutrient sharing with mucus-associated gut bacteria"

**Supplementary Tables:**

**Supplementary Table 1:** Enzyme names and cloning primers
**Supplementary Table 2:** Kinetics parameters of *A. muciniphila* fucosidases
**Supplementary Table 3:** Number of assigned *O*- and *N*-glycan structures studied in this work
**Supplementary Table 4:** Fucosidase relative activity towards porcine gastric, colonic mucin and fetuin
**Supplementary Table 5:** Normalized activity of *A. muciniphila* fucosidases on HMOs, mucins and fetuin
**Supplementary Table 6:** Activity profiles of *A. muciniphila* sialidases on porcine colonic mucin *O*-glycans
**Supplementary Table 7:** Normalised activities of *A. muciniphila* sialidases on HMOs and attached *O*-glycans from mucin and fetuin
**Supplementary Table 8:** The top structural orthologues of *Am*GH29D
**Supplementary Table 9:** The top structural orthologues of *Am*GHxxx
**Supplementary Table 10:** The top structural orthologues of the CBM-like domain of *Am*GHxxx
**Supplementary Table 11:** Inhibition of *A. muciniphila* fucosidases by 1-Deoxyfuconojirimycin (DFJ)
**Supplementary Table 12:** Inhibition of *A. muciniphila* sialidases by 2,3-dehydro-2-deoxy-*N*-acetylneuraminic acid (DANA)
**Supplementary Table 13:** The effect of fucosidase and sialidase inhibition on growth of *A. muciniphila* on PCM
**Supplementary Table 14:** Prevalence of fucosidase and sialidase genes in *A. muciniphila* genomes
**Supplementary Table 15:** Data collection and refinement statistics for *Am*GH29D
**Supplementary Table 16:** Data collection and refinement statistics for *Am*GHxxx

**Supplementary Figures:**

**Supplementary Fig. 1:** Modular organisation and sequence comparison of the *A. muciniphila* GH29 and GH95
**Supplementary Fig. 2:** Modular organisation of characterised GH29 and GH95 fucosidases
**Supplementary Fig. 3:** Phylogenetic clustering of *A. muciniphila* GH29 and GH95 fucosidases
**Supplementary Fig. 4:** Activity of *A. muciniphila* fucosidases on model oligosaccharides
**Supplementary Fig. 5:** Activity profiles of *A. muciniphila* fucosidases on selected mucin *N*- and *O*-glycans
**Supplementary Fig. 6:** Fucosidase activity on defined *O*-glycoprotein conjugated Lewis epitopes
**Supplementary Fig. 7:** Modular organization and phylogeny of *A. muciniphila* sialidases
**Supplementary Fig. 8:** Sialidase activity on oligosaccharides
**Supplementary Fig. 9:** Sialidase activity on mucin sialylated *O*-glycan motifs and immunoglobulin G *N*-glycans
**Supplementary Fig. 10:** Cobra strike pose architecture and putative binding domains of *Am*GH29D
**Supplementary Fig. 11:** Comparison of the ALPHA fold model of *Am*GH29C and the crystal structure of *Am*GH29D
**Supplementary Fig. 12:** Phylogenetic tree of GHxxx as well as active site and surface binding site motif conservation
**Supplementary Fig. 13:** Architecture of *Am*GHxxx and comparison to GH33 sialidases
**Supplementary Fig. 14:** Catalytic site signatures of *Am*GHxxx as compared to the closest GH33 sialidase
**Supplementary Fig. 15:** The NMR analysis of the inverting mechanism of *Am*GHxxx
**Supplementary Fig. 16:** Ligand binding at the active site and secondary surface binding sites of *Am*GHxxx
**Supplementary Fig. 17:** Binding of *A. muciniphila* fucosidases and sialidases to mucin
**Supplementary Fig. 18:** Localisation of fucosidase and sialidase activities of *A. muciniphila*
**Supplementary Fig. 19:** Growth of butyrate producing Lachnospiraceae on monosaccharides

**Supplementary Table 1: Enzyme names and cloning primers.**

| Locus tag | SP[a] | Enzyme | GenBank | Sense primer | Antisense primer |
|---|---|---|---|---|---|
| Amuc_0010 | 20 | *Am*GH29A | ACD03857.1 | **AGGAGATATACCATG**CAGTCC GCCACTAAAATCATTACG | **GGTGGTGGTGCTCGA**GTTTGCTCAGTT TGATGACGGAAC |
| Amuc_0146 | 23 | *Am*GH29B | ACD03990.1 | **AGGAGATATACCATG**GGGAAT GCCATCACCGTCC | **GGTGGTGGTGCTCGA**GTTGAAGTTTG ATGACGGTATCCAGC |
| Amuc_0392 | 36 | *Am*GH29C | ACD04231.1 | **AGGAGATATACCATG**GGCTGG ACCGCAGCACC | **GGTGGTGGTGCTCGA**GTTTGCCTGCG GGAGTGC |
| Amuc_0846 | 24 | *Am*GH29D | ACD04679.1 | **AGGAGATATACCATG**GGGCCG AAGGGCTGTTTAAC | **GGTGGTGGTGCTCGA**GTTTTCCCAATA CGCCCAGCTC |
| Amuc_0186 | 23 | *Am*GH95A | ACD04030.1 | **AGGAGATATACCATG**GCCATT CCGGCCCCCATG | **GGTGGTGGTGCTCGA**GATGGGAAAGC GGAGGAAAATCAAG |
| Amuc_1120 | 13 | *Am*GH95B | ACD04946.1 | **AGGAGATATACCATG**AGTGCC GTTTCTTTCGGGTGG | **GGTGGTGGTGCTCGA**GCCTGGCCGCG GGCTG |
| Amuc_0623 | 21 | | ACD04460.1 | **AGGAGATATACCATG**ACCGTA CCGGCCCATTCC | **GGTGGTGGTGCTCGA**GGGGACGTTTC AGAAGGCGATTAAC |
| Amuc_0625 | 38 | *Am*GH33A | ACD04462.1 | **AGGAGATATACCATG**CAGGAA GAGAAAACCGGTTTCC | **GGTGGTGGTGCTCGA**GCTTGAGAACA GGAGCTTTTTTGC |
| Amuc_1547 | 22 | *Am*GH33B | ACD05368.1 | **AGGAGATATACCATG**GCACCC GTTCCGGAAC | **GGTGGTGGTGCTCGA**GCTTCACCCGG GCATTCAC |
| Amuc_1835 | 17 | *Am*GH33C | ACD05653.1 | **AGGAGATATACCATG**GGCAAG GAAAGCTTTGAGCAGG | **GGTGGTGGTGCTCGA**GGCGCGCATTT TTGCGTTAAG |

[a]The size of the signal peptide in amino acids as predicted from SignalP (V.5.0) (see Materials and methods section).
The cloning vector homologous recombination patches to the cloning cassette of the used pET28a(+) vector are in bold. The *A. muciniphila* sialidases with the locus tags Amuc_0623, Amuc_0625, Amuc_1547 and Amuc_1835 have been referred to as Am0705, Am0707, Am1757 and Am2085 respectively[1], in previous work that used fluorescently labelled model substrates to indirectly show sialidase activity.

1. Huang K*, et al.* Biochemical characterization of the neuraminidase pool of the human gut symbiont *Akkermansia muciniphila*. *Carbohydr Res* **415**, 60-65 (2015).

**Supplementary Table 2: Kinetics parameters of *A. muciniphila* fucosidases.**

| | Enzyme | | | | | |
|---|---|---|---|---|---|---|
| | ***Am*GH29A** | ***Am*GH29B** | ***Am*GH29C** | ***Am*GH29D** | ***Am*GH95A** | ***Am*GH95B** |
| $k_{cat}$ ($min^{-1}$) | 2610 ± 90 | 6.22 ± 0.40 | 0.60 ± 0.01 | 1.29 ± 0.06 | 58.7 ± 1.01 | 22.2 ± 1.02 |
| $K_M$ (mM) | 1.06 ± 0.11 | 1.63 ± 0.33 | 4.12 ± 0.41 | 2.52 ±0.34 | 0.76 ± 0.04 | 2.98 ± 0.46 |
| $k_{cat}$ /$K_M$ ($min^{-1}$·$mM^{-1}$) | 2.470 ± 90 | 3.81 ± 0.23 | 8.70 ± 0.01 | 0.57 ± 0.03 | 77.5 ± 1.34 | 7.46 ± 0.34 |

The kinetic parameters were determined towards the model substrate *para*-nitrophenyl-α-L-Fucoside (*p*NPFuc) at 37 °C in 20 mM HEPES, 150 mM NaCl, pH 6.8. Data are mean values of three independent experiments (n=3) with standard deviation (SD).

**Supplementary Table 3: Number of assigned *O*- and *N*-glycan structures studied in this work.**

| Substrate | Number of assigned glycan structures in the fucosidase analysis | | |
|---|---|---|---|
| | Total | Fucosylated | Sialylated |
| *O*-glycans from PGM, PCM and fetuin | 160 | 88 | 44 |
| *N*-glycans from human IgG | 22 | 15 | 10 |
| **Substrate** | **Number of assigned glycan structures in the sialidase analysis** | | |
| | Total | Fucosylated | Sialylated |
| *O*-glycans from PCM | 80 | 41 | 30 |
| *O*-glycans from $Muc2_{Mouse}$ | 74 | 17 | 36 |
| *N*-glycans from human IgG | 24* | 13 | 10 |

Number of glycan structures that are assigned from the LC-MS analysis of reactions of *A. muciniphila* fucosidases and sialidases on different substrates. *The glycans are from a different experiments than the fucosidases.

**Supplementary Table 4: Fucosidase relative activity towards porcine gastric, colonic mucin and fetuin.**

| | *O*-glycans | | | |
|---|---|---|---|---|
| | **non-fucosylated** | **Fuc-α1,2** | **Fuc-α1,2 & α1,3/4** | **Fuc-α1,3/4** |
| **Enzyme** | (%) | (%) | (%) | (%) |
| Control | 57 | 22 | 11 | 11 |
| *Am*GH29A | 52 | 25 | 11 | 12 |
| *Am*GH29B | 56 | 23 | 9 | 12 |
| *Am*GH29C | 69 | 30 | 0 | 0 |
| *Am*GH29D | 60 | 30 | 4 | 6 |
| *Am*GH95A | 59 | 19 | 9 | 13 |
| *Am*GH95B | 74 | 3 | 0 | 23 |

Abundance (%) of non-fucosylated and fucosylated *O*-glycans present in a 1:1:1 per weight mixture of porcine gastric mucin, porcine colonic mucin and fetuin before and after incubation with *A. muciniphila* fucosidases or with buffer as control. Abundances are calculated based on relative intensities of 166 assigned individual glycan structures detected by LC-MS. Data are from a single (n=1) experiment.

**Supplementary Table 5: Normalized activity of *A. muciniphila* fucosidases on HMOs, mucins and fetuin.**

| Substrate | Enzyme | | | | | |
|---|---|---|---|---|---|---|
| | ***Am*GH29A** ($min^{-1}$) | ***Am*GH29B** ($min^{-1}$) | ***Am*GH29C** ($min^{-1}$) | ***Am*GH29D** ($min^{-1}$) | ***Am*GH95A** ($min^{-1}$) | ***Am*GH95B** ($min^{-1}$) |
| HMOs | 37.5 ± 5.52 | 5.04 ± 0.63 | 263 ± 2.57 | 239 ± 1.55 | 232 ± 4.68 | 238 ± 7.72 |
| Fetuin | N.D. | N.D. | N.D. | N.D. | N.D. | N.D. |
| PGM | 1.65 ± 0.10 | 1.14 ± 0.09 | 3.53 ± 0.03 | 2.69 ± 0.09 | 21.7 ±0.24 | 268 ± 5.08 |
| PCM | 0.35 ± 0.00 | 0.39 ± 0.01 | 67.0 ± 0.01 | 27.4 ± 1.10 | 5.67 ± 0.82 | 47.6 ± 2.80 |

Normalized activity (*V*/E) determined towards 0.5 % (w/v) substrate concentration and a with an enzyme concentration= 0.5 µM. Enzymatic reactions were performed for 1 h, except for reactions containing fetuin which were incubated for 3 h. N.D. not detected. The very low activity of *Am*GH29A and *Am*GH29B on HMOs is likely attributed to activity on the trisaccharide 2'-fucosyl lactose (2FL, see Supplementary Fig. 3j). Data are mean values of three independent experiments (n=3) with standard deviation (SD).

**Supplementary Table 6: Activity profiles of *A. muciniphila* sialidases on porcine colonic mucin *O*-glycans.**

| | ***O*-glycans** | | | |
|---|---|---|---|---|
| **Enzyme** | **non-sialylated** (%) | **α2,3** (%) | **α2,3 & α2,6** (%) | **α2,6** (%) |
| Control | 72 | 7 | 2 | 20 |
| Amuc_0623 | 75 | 6 | 2 | 17 |
| *Am*GH33A | 97 | 3 | 0 | 0 |
| *Am*GH33B | 99 | 1 | 0 | 0 |
| *Am*GHxxx | 76 | 3 | 0 | 21 |

Abundance (%) of non-sialylated and differentially sialylated *O*-glycans from porcine colonic mucin before and after incubation with *A. muciniphila* sialidases and or with buffer as control. Abundances are calculated based on relative intensities of 82 assigned individual glycan structures detected by LC-MS. The data are for both Neu5Ac and Neu5Gc forms of sialic acid that are cleaved by the enzymes. Data are from a single (n=1) experiment.

**Supplementary Table 7: Normalised activities of *A. muciniphila* sialidases on HMOs and attached *O*-glycans from mucin and fetuin.**

| | Enzyme | | | |
|---|---|---|---|---|
| **Substrate** | **Amuc_0623** (min$^{-1}$) | ***Am*GH33A** (min$^{-1}$) | ***Am*GHxxx** (min$^{-1}$) | ***Am*GH33B** (min$^{-1}$) |
| HMOs | 0.20 ± 0.03 | 9.98 ± 0.88 | 0.76 ± 0.18 | 10.8 ± 1.30 |
| Fetuin | 2.10 ± 0.36 | 9.14 ± 1.21 | 4.53 ± 2.16 | 12.5 ± 2.97 |
| PGM | N.D. | 0.86 ± 0.27 | N.D. | 2.21 ± 0.37 |
| PCM | 0.35 ± 0.13 | 1.15 ± 0.27 | 0.81 ± 0.18 | 2.19 ± 0.25 |

Normalized activity (*V*/E) determined towards 0.5 % (w/v) substrate concentration and with an enzyme concentration= 0.5 µM. N.D. not detected within 1 h or 3 h assays. Activity of *Am*GHXXX on HMOs likely attributed to the very low activity on 3’-Sialyl lactose. Data are means of triplicates (n=3) with standard deviation (SD).

**Supplementary Table 8: The top structural orthologues of *Am*GH29D.**

| | PDB ID | Z-score | RMSD (Å) | Aligned[a] | Total[b] | Identity (%) | name/GH/Source organism |
|---|---|---|---|---|---|---|---|
| 1 | 6OR4-B | 50.9 | 1.5 | 438 | 449 | 39 | SpGH29/GH29/*Streptococcus pneumoniae* TIGR4 |
| 2 | 5K9H-A | 50.3 | 7.5 | 457 | 554 | 39 | GH29_0940/GH29/Rumen unknown bacteria |
| 3 | 3UES-A | 48.8 | 1.5 | 433 | 457 | 41 | BiAfcB/GH29/*Bifidobacterium longum* subsp. infantis ATCC 15697 |
| 4 | 4OZO-B | 48.1 | 1.8 | 439 | 459 | 40 | BT2192/GH29/*Bacteroides thetaiotaomicron* VPI-5482 |
| 5 | 4zrx-A | 45.5 | 3.0 | 451 | 581 | 42 | Bovatus_01698/GH29/ *Bacteroides ovatus* ATCC 8483 |
| 6 | 6tr3-A | 42.6 | 2.2 | 448 | 505 | 34 | CDL26_02305/GH29/*Ruminococcus gnavus* GH29 fucosidase E1 |
| 7 | 3gza-b | 41.5 | 1.9 | 396 | 431 | 30 | BT3798/GH29/*Bacteroides thetaiotaomicron* VPI-5482 |
| 8 | 6gn6-C | 33.5 | 2.8 | 301 | 421 | 22 | aLfuk1/GH29/*Paenibacillus thiaminolyticus* |
| 9 | 6o1j-A | 32.6 | 2.0 | 272 | 329 | 25 | BN194_28780/GH29/*Lacticaseibacillus casei* W56 |
| 10 | 4jfs-B | 31.4 | 2.7 | 307 | 437 | 26 | BtFuc2970/GH29/ |

The data are based on a DALI search and only the top hit of mutants and/or complexes with ligands is included to avoid redundancy. [a]Aligned residues between the hit protein and *Am*GH29D. [b]Total number of residues in the hit protein.

**Supplementary Table 9: The top structural orthologues of *Am*GHxxx.**

| | PDB ID | Z-score | RMSD (Å) | Aligned[a] | Total[b] | Identity (%) | name/GH/Source organism |
|---|---|---|---|---|---|---|---|
| 1 | 1W8O-A | 26.2 | 3.6 | 311 | 601 | 16 | NedA sialidase/GH33 *Micromonospora viridifaciens* |
| 2 | 1SNT-A | 26.1 | 2.8 | 281 | 352 | 10 | Neu2 sialidase/GH33 *Homo sapien* |
| 3 | 2VK7-B | 25.6 | 2.9 | 283 | 448 | 14 | NanI sialidase/GH33 *Clostridium perfringens* |
| 4 | 3H73-B | 25.4 | 3.0 | 286 | 477 | 16 | NanA sialidase/GH33 *Streptococcus pneumoniae* |
| 5 | 5HX0-B | 24.4 | 3.0 | 279 | 364 | 24 | Unknown protein/GH33 Dyadobacter fermentans |
| 6 | 4XJZ-A | 23.9 | 3.1 | 280 | 658 | 20 | NanB sialidase/GH33/ *Streptococcus pneumoniae* |
| 7 | 2SLI-A | 23.8 | 3.0 | 267 | 679 | 18 | Intramolecular transsialidase L (MDSA)/GH33/*Macrobdella decora* |
| 8 | 4X47-A | 23.5 | 3.0 | 275 | 489 | 20 | Anhydrosialidase RgNanH/GH33/*Ruminococcus gnavus* |
| 9 | 6MYV-A | 23.3 | 3.0 | 277 | 522 | 20 | Sialidase/GH33/unidentified bacterium |
| 10 | 4YZ2-A | 23.2 | 2.9 | 272 | 655 | 19 | NanC sialidase/GH33 *Streptococcus pneumoniae* |

Based on a DALI search; only the top hit of mutants and/or complexes with ligands is included to avoid redundancy. [a]Aligned residues between the hit protein and AmGH33B. [b]Total number of residues in the hit protein.

**Supplementary Table 10: The top structural orthologues of the CBM-like domain of *Am*GHxxx.**

| | PDB ID | Z-score | RMSD (Å) | Aligned[a] | Total[b] | Identity (%) | name/GH/Source organism |
|---|---|---|---|---|---|---|---|
| 1 | 5MQR-A | 15.4 | 2.1 | 127 | 1082 | 15 | BT1020/GH33/*Bacteroides thetaiotaomicron ATCC 29148* |
| 2 | 4AGG-A | 12.9 | 2.8 | 121 | 144 | 11 | Galectin/unknown *Cinachyrelle* sp./marine sponge |
| 3 | 4HLO-A | 12.6 | 2.2 | 114 | 278 | 7 | Galectin/*Tocascaris leonine*/helminth parasite |
| 4 | 3AFK-A | 12.5 | 2.8 | 128 | 168 | 11 | Galectin/*Cyclocybe aegerita*/fungi |
| 5 | 5XRM-A | 12.3 | 2.6 | 117 | 141 | 18 | Galectin/Homo sapiens |
| 6 | 2A6Y-A | 11.5 | 3.1 | 140 | 231 | 9 | Carbohydrate recognition domain/*Saccharomyces cerevisiae* |
| 7 | 2WSU-B | 11 | 2.5 | 117 | 306 | 10 | Galectin/ porine adenovirus 4 |
| 8 | 5N8K-A | 10.7 | 2.6 | 123 | 644 | 11 | β-galactocerebrosidase/*Mus musculus* |
| 9 | 1KIT-A | 10.7 | 2.6 | 119 | 757 | 11 | GH33/*Vibrio cholerae* |
| 10 | 3WUC-B | 10.7 | 2.6 | 113 | 137 | 14 | Galectin/*Xenopus laevis* |

The data are based on a DALI search and only the top hit of mutants and/or complexes with ligands is included to avoid redundancy. [a]Aligned residues between the hit protein and CBM-like domain of *Am*GHxxx. [b]Total number of residues in the hit protein.

**Supplementary Table 11: Inhibition of *A. muciniphila* fucosidases by 1-Deoxyfuconojirimycin (DFJ).**

| | Enzyme | | | | | |
|---|---|---|---|---|---|---|
| | ***Am*GH29A** | ***Am*GH29B** | ***Am*GH29C** | ***Am*GH29D** | ***Am*GH95A** | ***Am*GH95B** |
| $IC_{50}$ (µM) | 0.60 ± 0.02 | 1.50 ± 0.10 | 13.7 ± 0.41 | 7.29 ± 0.16 | 53.7 ± 5.9 | 24.0 ± 1.6 |

The inhibition constant $IC_{50}$ was determined towards 2 mM *p*NPFuc substrate concentration with inhibitor concentrations in the 0.1 - 100 µM range. The enzyme concentration=0.5 µM for all fucosidases except for *Am*GH29C and *Am*GH29D, which were assayed at concentration= 10 µM due to their low activity. Data are the means of three independent experiments (n=3) with standard deviation (SD)

**Supplementary Table 12: Inhibition of *A. muciniphila* sialidases by 2,3-dehydro-2-deoxy-*N*-acetylneuraminic acid (DANA).**

| | Enzyme | | | |
|---|---|---|---|---|
| | **Amuc_0623** | ***Am*GH33A** | ***Am*GH33B** | ***Am*GHxxx** |
| $IC_{50}$ (µM) | N.D. | 61.4 ± 2.20 | 133 ± 11.1 | 199 ± 13.2 |
| Normalized activity (Emission units $min^{-1}$ $nM^{-1}$) | 0.22 ± 0.04x$10^{-1}$ | 6.50 ± 0.41 | 4.46 ± 0.13 | 2.18 ± 0.10 |

$IC_{50}$ constants determined towards 1 mM 4-Methylumbelliferyl *N*-acetyl-a-D-neuraminic acid sodium salt (4MU-Neu5Ac) with an inhibitor concentration in the 0.01-1 mM range. The enzyme concentration was 50 nM for all sialidases, except Amuc_0623 which was assayed at 200 nM. Enzymatic reactions were performed for 30 min. N.D. low affinity towards DANA reflected by the lack of curvature, which precluded reliable determination of inhibition constant. The normalized activity, expression in arbitrary emission units per min and nM enzyme is also shown, to depict that the enzymes have different activity levels on this substrate. Data are means of three independent experiments (n=3) with standard deviation (SD).

**Supplementary Table 13: The effects of fucosidase and sialidase inhibition on growth of *A. muciniphila* on PCM.**

| Time (h) | | Growth substrate | | | |
|---|---|---|---|---|---|
| | | PCM | PCM + Inhibitors | GlcNAc+GalNAc | GlcNAc+GalNAc+ Inhibitors |
| 5[a] | $OD_{600}$ | 0.41 ± 0.01 | 0.14 ± 0.01 | 0.21 ± 0.01 | 0.20 ± 0.01 |
| | *p*-value | $P < 1.1 \times 10^{-8}$ | | P < 0.62 | |
| 8[a] | $OD_{600}$ | 1.11 ± 0.04 | 0.19 ± 0.02 | 0.39 ± 0.02 | 0.40 ± 0.05 |
| | *p*-value | $P < 8.59 \times 10^{-9}$ | | P < 0.58 | |
| 24[a] | $OD_{600}$ | 1.39 ± 0.04 | 0.46 ± 0.03 | 1.32 ± 0.02 | 1.32 ± 0.04 |
| | *p*-value | $P < 1.42 \times 10^{-8}$ | | P < 0.93 | |
| 24[b] | $OD_{600}$ | 1.41 ± 0.02 | 0.03 ± 0.01 | 1.43 ± 0.06 | 1.45 ± 0.05 |
| | *p*-value | $P < 4.45 \times 10^{-11}$ | | P < 0.72 | |

Growth level of *A. muciniphila* in the absence or presence of an equimolar fucosidase and sialidase inhibitor blend on PCM or an equimolar mixture of GlcNAc/GalNAc after 5, 8 and 24h. Growth media were supplemented with 0.5% (w/v) carbohydrates and DFJ/DANA to a final concentration of 1 mM[a] or 20[b] mM each. The growth experiments were performed in 4 independent biological replicates (n=4) and the data are presented as mean values with standard deviations.

**Supplementary Table 14: Prevalence of fucosidase and sialidase genes in *A. muciniphila* genomes**

| | Enzyme | | | | | |
|---|---|---|---|---|---|---|
| Fucosidase | *Am*GH95A | *Am*GH95B | *Am*GH29A | *Am*GH29B | *Am*GH29C | *Am*GH29D |
| Prevalence (%) | 93.8 | 98.5 | 99.4 | 37.3 | 99.5 | 89.9 |
| Sialidase | Amuc_0623 | *Am*GH33A | *Am*GH33B | *Am*GHxxx | | |
| Prevalence (%) | 24.4 | 98.3 | 98.9 | 98.3 | | |

Global prevalence of fucosidases and sialidases in 177 *A. muciniphila* genomes of human origin (see materials and methods). The prevalence of fucosidases and sialidases genes were analyzed by a BLASTP search using the amino acid sequences of *Am*GH95A, *Am*GH95B, *Am*GH29A, *Am*GH29B, *Am*GH29C, *Am*GH29D, Amuc_0623, *Am*GH33A, *Am*GH33B and *Am*GHxxx from *Akkermansia muciniphila* ATCC BAA-835 (same as *Akkermansia muciniphila* DSM 22959) as query.

| Time (h) | | Growth substrate | | | |
|---|---|---|---|---|---|
| | | PCM | PCM + Inhibitors | GlcNAc/GalNAc | GlcNAc/GalNAc + Inhibitors |
| 5 | *OD600* | 0.41 ± 0.01 | 0.14 ± 0.01 | 0.21 ± 0.01 | 0.20 ± 0.01 |
| | *p-value* | p < 1.1 x 10-8 | | *p* < 0.62 | |
| 8 | *OD600* | 1.11 ± 0.04 | 0.19 ± 0.02 | 0.39 ± 0.02 | 0.40 ± 0.05 |
| | *p-value* | p < 8.59 x 10-9 | | *p* < 0.58 | |
| 24 | *OD600* | 1.39 ± 0.04 | 0.46 ± 0.03 | 1.32 ± 0.02 | 1.32 ± 0.04 |
| | *p-value* | *p* < 1.42 x 10-8 | | *p* < 0.93 | |

**Supplementary Table 15: Data collection and refinement statistics of *Am*GH29D**

| | **No additives** |
|---|---|
| PDB accession | 8AYR |
| Resolution range(Å)* | 34.45 - 2.70 (2.798 - 2.70) |
| Space group | P 1 |
| Unit cell (Å, °) | 76.19 76.45 84.5 88.41 89.08 88.23 |
| Total reflections | 184329 (14917) |
| Unique reflections | 47292 (3963) |
| Multiplicity | 3.9 (3.8) |
| Completeness (%) | 90.33 (75.09) |
| Mean I/sigma(I) | 9.93 (1.63) |
| Wilson B-factor | 59.87 |
| R-merge | 0.082 (0.67) |
| R-meas | 0.094 (0.78) |
| R-pim | 0.048 (0.40) |
| CC1/2 | 0.99 (0.86) |
| CC* | 0.99 (0.96) |
| Reflections used in refinement | 47185 (3953) |
| Reflections used for R-free | 2450 (205) |
| R-work | 0.23 (0.42) |
| R-free | 0.27 (0.47) |
| CC(work) | 0.95 (0.84) |
| CC(free) | 0.92 (0.72) |
| Number of non-hydrogen atoms | 10626 |
| macromolecules | 10602 |
| ligands | 4 |
| solvent | 20 |
| Protein residues | 1353 |
| RMS(bonds, Å) | 0.005 |
| RMS(angles,°) | 0.86 |
| Ramachandran favoured (%) | 91.48 |
| Ramachandran allowed (%) | 8.01 |
| Ramachandran outliers (%) | 0.52 |
| Rotamer outliers (%) | 0.82 |
| Clashscore | 10.98 |
| Average B-factor ($Å^2$) | 90.90 |
| macromolecules | 90.94 |
| ligands | 96.59 |
| solvent | 68.08 |
| Number of TLS groups | 6 |

*Statistics for the highest-resolution shell are shown in parentheses

**Supplementary Table 16: Data collection and refinement statistics of *Am*GHxxx.**

| | No ligand | DANA | DANA and GNB |
|---|---|---|---|
| PDB accession | 8AXT | 8AXS | 8AXI |
| Resolution range(Å)* | 53.74 - 1.59 (1.647 - 1.59) | 46.35 - 1.3 (1.346 - 1.3) | 47.23 - 1.25 (1.295 - 1.25) |
| Space group | P 1 21 1 | P 1 21 1 | P 1 21 1 |
| Unit cell(Å, °) | 72.96 56.79 146.72 90 94.55 90 | 104.07 51.78 118.44 90 93.07 90 | 103.92 51.51 118.47 90 93.15 90 |
| Total reflections | 1073418 (104775) | 1692885 (126087) | 2015188 (138573) |
| Unique reflections | 160820 (15926) | 308437 (29552) | 323490 (25068) |
| Multiplicity | 6.7 (6.6) | 5.5 (4.2) | 6.2 (5.5) |
| Completeness (%) | 99.77 (99.70) | 99.38 (95.93) | 93.50 (72.90) |
| Mean I/sigma(I) | 6.66 (1.35) | 11.78 (0.87) | 15.68 (2.50) |
| Wilson B-factor | 15.12 | 16.59 | 14.13 |
| R-merge | 0.29 (>1) | 0.063 (>1) | 0.049 (0.63) |
| R-meas | 0.32 (>1) | 0.069 (>1) | 0.053 (0.70) |
| R-pim | 0.123(<1) | 0.027 (0.8868) | 0.021 (0.28) |
| CC1/2 | 0.99 (0.33) | 0.999 (0.48) | 0.999 (0.87) |
| CC* | 0.997 (0.71) | 1 (0.81) | 1 (0.96) |
| Reflections used in refinement | 160792 (15926) | 307572 (29515) | 323214 (25037) |
| Reflections used for R-free | 8029 (805) | 15493 (1486) | 16213 (1226) |
| R-work | 0.168 (0.29) | 0.155 (0.42) | 0.153 (0.25) |
| R-free | 0.202 (0.33) | 0.175 (0.43) | 0.169 (0.27) |
| CC(work) | 0.963 (0.75) | 0.975 (0.68) | 0.970 (0.92) |
| CC(free) | 0.948 (0.62) | 0.969 (0.67) | 0.965 (0.90) |
| Number of non-hydrogen atoms | 10349 | 10778 | 10756 |
| macromolecules | 9014 | 9212 | 9057 |
| ligands | 7 | 180 | 240 |
| solvent | 1328 | 1467 | 1573 |
| Protein residues | 1142 | 1145 | 1143 |
| RMS(bonds, Å) | 0.010 | 0.012 | 0.011 |
| RMS(angles, °) | 1.05 | 1.06 | 1.11 |
| Ramachandran favoured (%) | 96.40 | 96.49 | 96.75 |
| Ramachandran allowed (%) | 3.43 | 3.34 | 3.25 |
| Ramachandran outliers (%) | 0.18 | 0.18 | 0.00 |
| Rotamer outliers (%) | 0.31 | 0.41 | 0.31 |
| Clashscore | 2.40 | 2.54 | 3.07 |
| Average B-factor($Å^2$) | 19.75 | 24.20 | 21.80 |
| macromolecules | 18.48 | 22.67 | 19.99 |
| ligands | 21.65 | 31.64 | 32.39 |
| solvent | 28.36 | 33.29 | 31.36 |
| Number of TLS groups | 11 | 12 | 11 |

*Statistics for the highest-resolution shell are shown in parentheses.

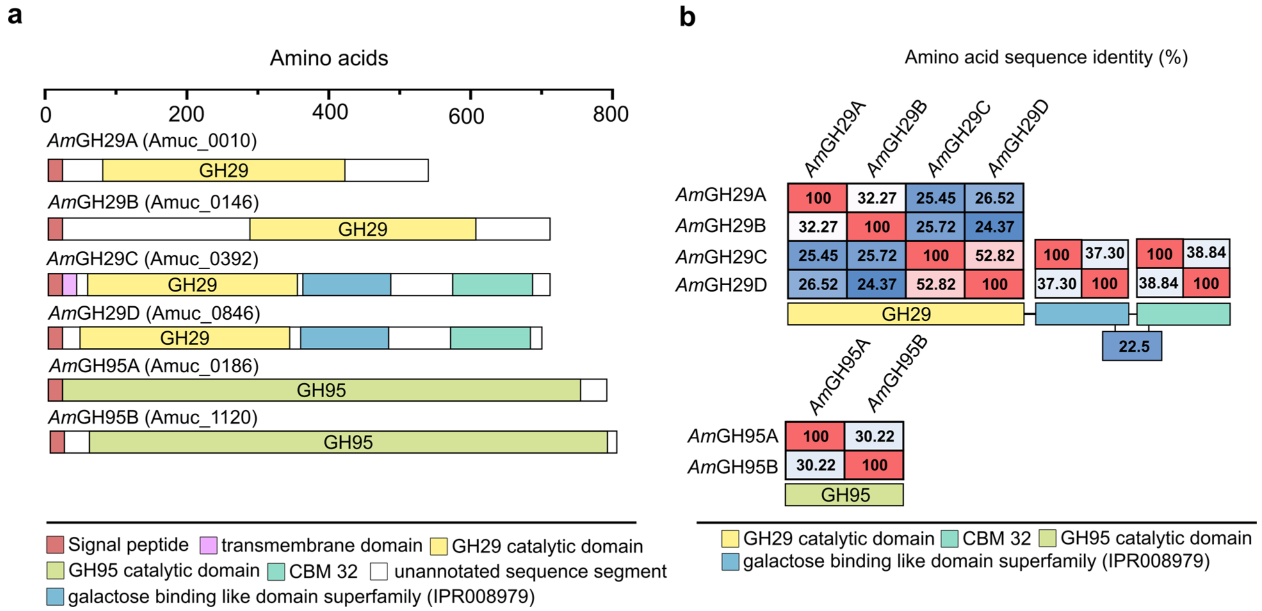


**Supplementary Fig. 1: Modular organization and sequence comparison of *A. muciniphila* GH29 and GH95. a,** Size and modular organization based on annotations by CAZy, dbCAN meta server, InterPro, and signal peptide predictions using SignalP (v.5.0). **b,** Amino acid sequence identity matrix showing the evolutionary relationships amongst the catalytic modules of *A. muciniphila* GH95s and GH29s and well as within the putative CBMs of the *A. muciniphila* GH29s.

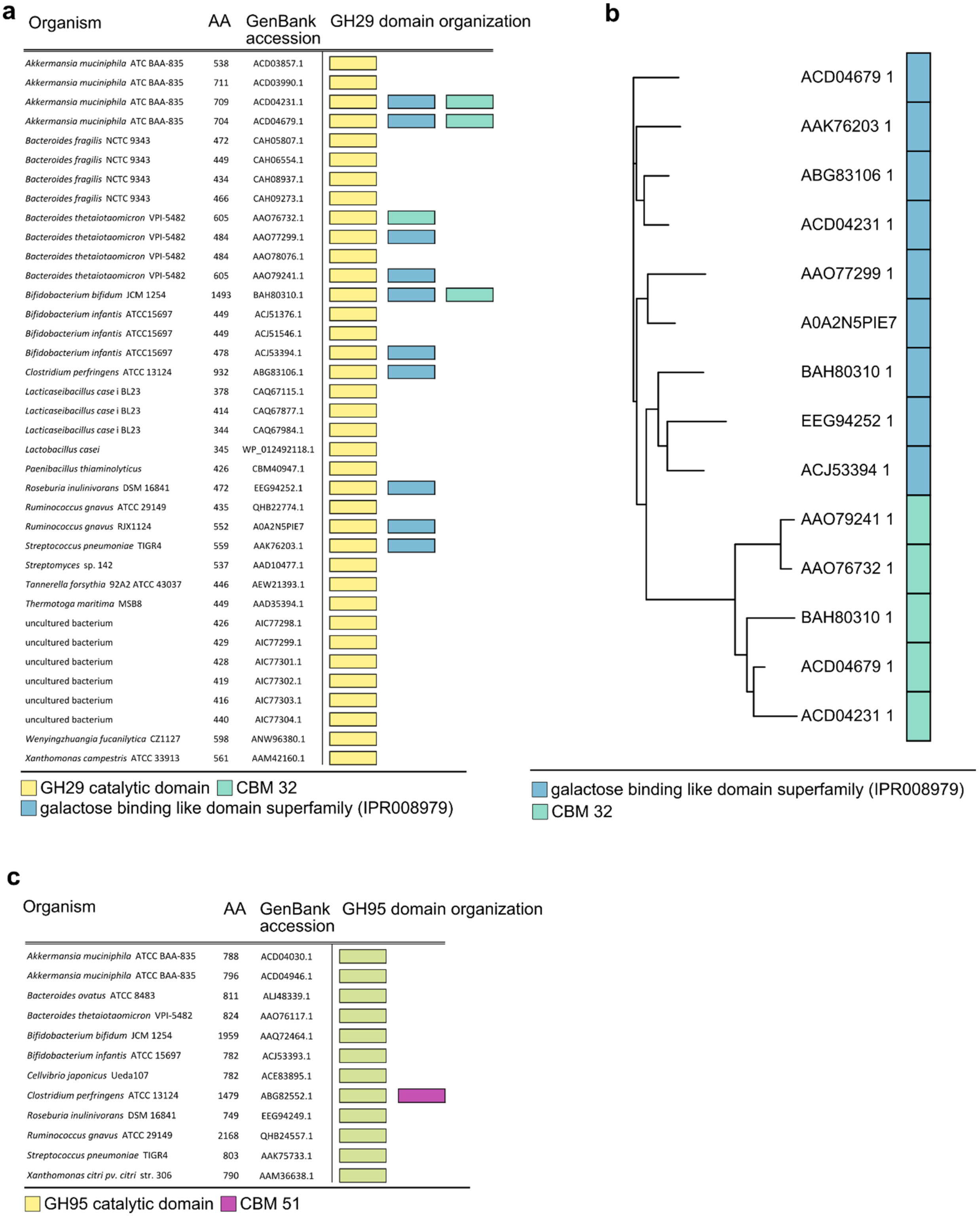


**Supplementary Fig. 2: Modular organization of characterized GH29 and GH95 fucosidases. a,** Size and domain organization of the four *A. muciniphila* GH29s and of previously characterized GH29 enzymes. **b,** Phylogenetic analysis showing the segregation of the galactose-binding-like domains and the CBM32 that occur in previously characterized GH29s and in *A. muciniphila* GH29 enzymes described in the present study. **c.** Size and modular organization of the two *A. muciniphila* GH95 fucosidases and of hitherto characterized GH95 enzymes. Domain annotations are from CAZy, dbCAN meta server and InterPro.

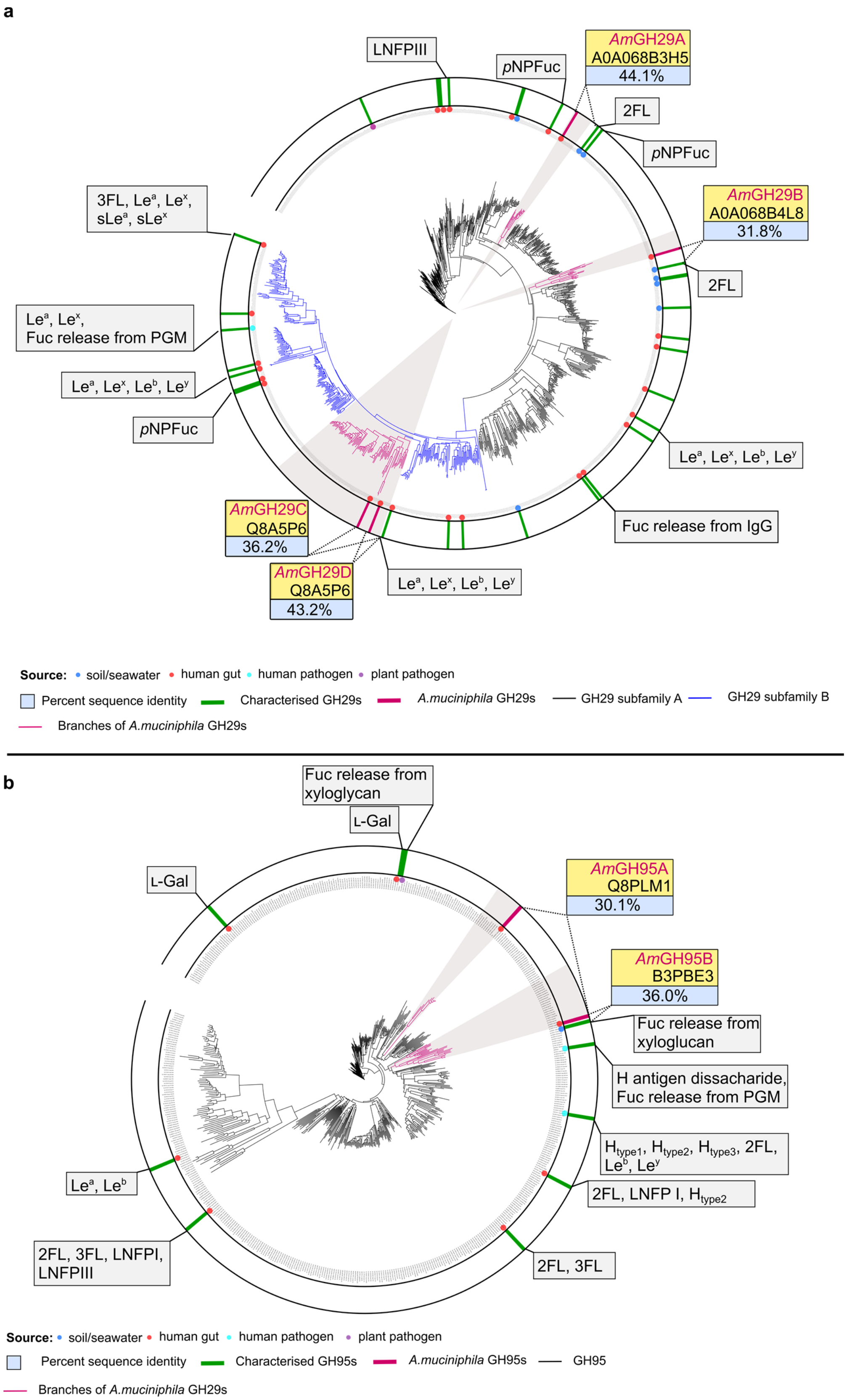


**Supplementary Fig. 3: Phylogenetic clustering of the *A. muciniphila* GH29 and GH95 fucosidases. a,** Phylogenetic tree of 1117 putative GH29 sequences. **b,** Phylogenetic tree of 543 putative GH95 sequences. Characterized GH29s and GH95s (as defined in CAZy) are green stripes, *A. muciniphila* GH29s and GH95s are in magenta stripes. *A. muciniphila* fucosidases and their closest characterized orthologues (indicated with their UniProt IDs) are in yellow boxes and the amino acid sequence identities between their catalytic modules are in the blue box. The source niches of the described fucosidases are indicated by coloured circles and the substrates the enzymes have been shown to be active on are shown. The sequences belonging to GH29 subfamily A (high activity on *p*NPFuc) and B (low activity on *p*NPFuc) are in black and blue, respectively. The branches populated by the *A. muciniphila* enzymes are in pink and highlighted by a grey shadow.

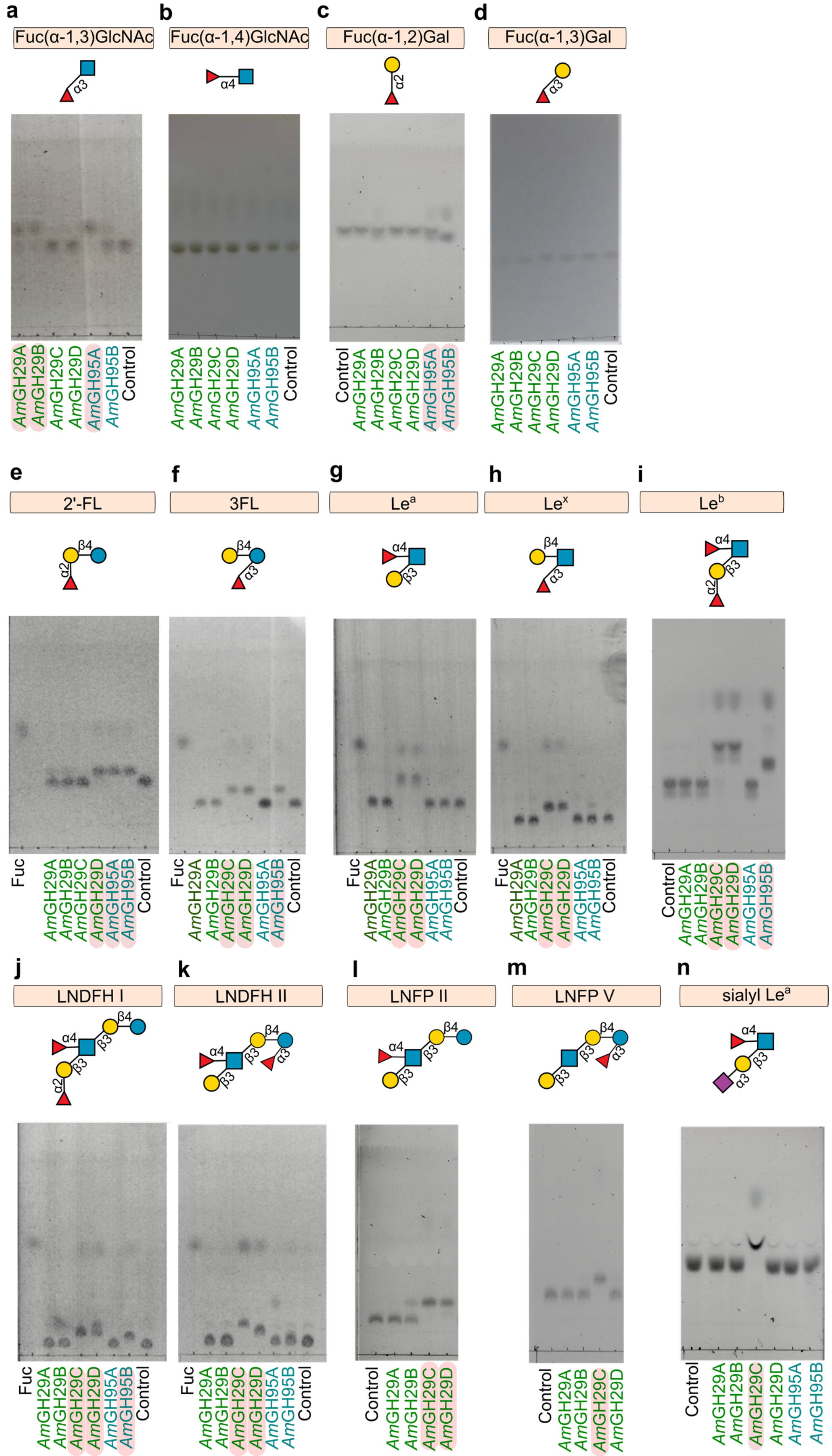


**Supplementary Fig. 4: The activity of *A. muciniphila* fucosidases on model oligosaccharides. a-n,** Fucosidase activity analysed using thin layer chromatography on di- and oligosaccharides. The enzymes that display activity are highlighted by a pink box. Source data are provided as a Source Data file labelled with the corresponding figure number and panel definition. The experiments were performed in 3 independent experiments (n=3), whereby the triplicate analyses yielded similar results.

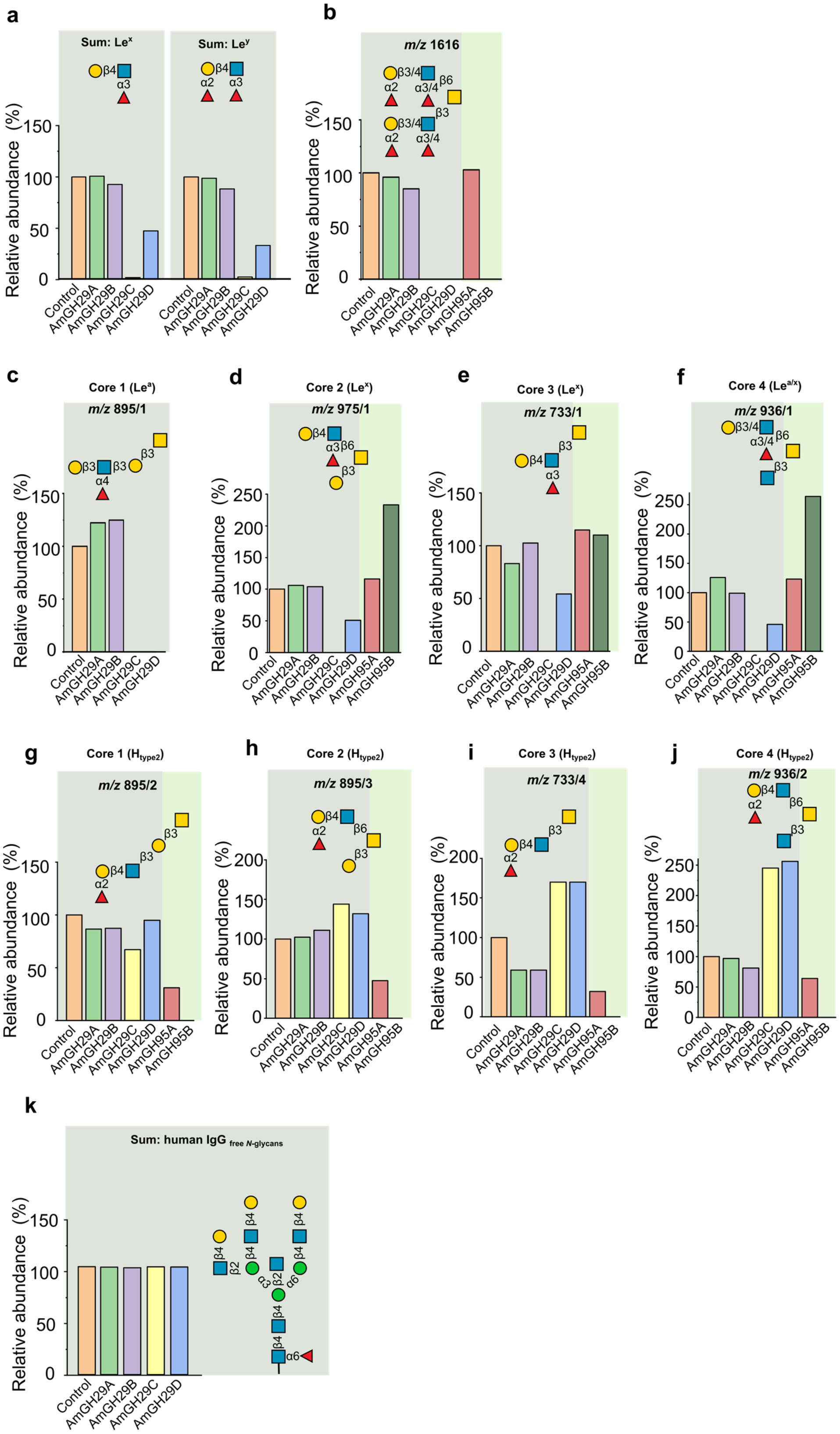

**Supplementary Fig. 5: Activity profiles of *A. muciniphila* fucosidases on selected mucin *O*-glycans and *N*-glycans.** The selected *O*-glycans are from intact colonic mucin, whereas the *N*-glycans were enzymatically released prior to the analysis **a,** Activity profiles showing the tolerance of adjacent double fucosylation. **b,** Activity profiles showing the tolerance of double branch fucosylations. **c,** Activity profiles showing the tolerance on the Le$^{a}$ epitope on core 1 structures. **d,** Activity profiles of *Am*GH29C and *Am*GH29D on core 2 structures. **e,** Activity profiles of *Am*GH29C and *Am*GH29D on core 3 structures. **f,** Activity profiles of *Am*GH29C and *Am*GH29D on core 4 structures. **g,** Activity profiles of *Am*GH95A and *Am*GH95B on core 1 structures. **h,** Activity profiles of *Am*GH95A and *Am*GH95B on core 2. **i,** Activity profiles of *Am*GH95A and *Am*GH95B on core 3 structures. **j.** Activity profiles of *Am*GH95A and *Am*GH95B on core 3 structures. **k,** Profiles showing lack of activity towards α1,6 linked fucosyl, present in human *N*-glycans. For isobaric glycans structures, the number after the slash (/n) denotes the corresponding structure in the LC-MS data file (see extended data). Data are from a single (n=1) experiment.

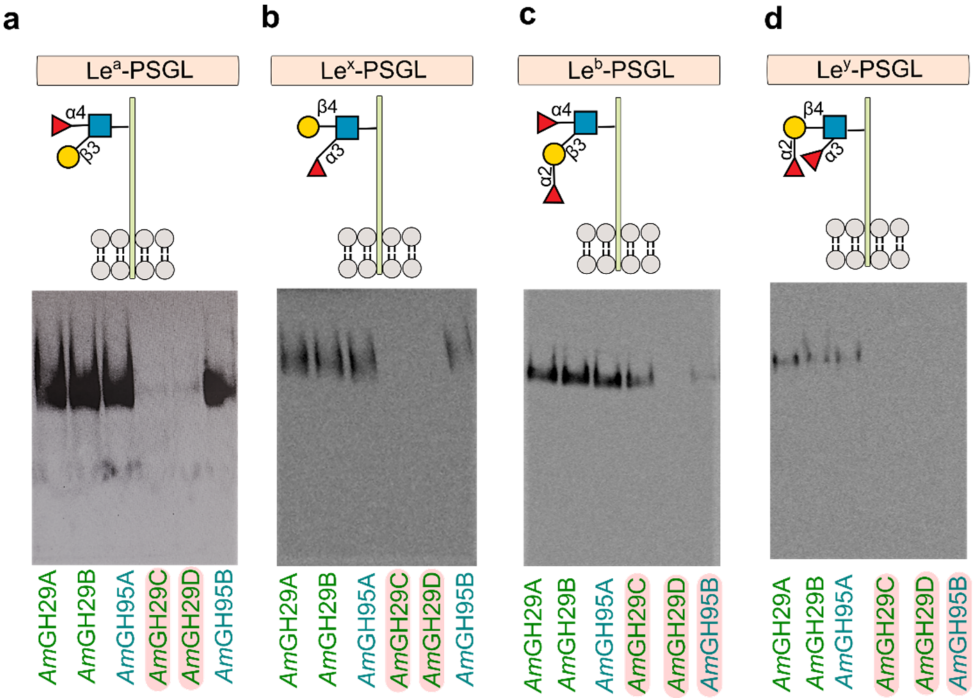


**Supplementary Fig. 6: Fucosidase activity on defined *O*-glycoprotein conjugated Lewis epitopes. a-d,** The fucosidase activity, which is observed as a decrease/loss of Western blot signal, on defined conjugated Lewis epitopes presented by the recombinant glyco-engineered P-selectin glycoprotein ligand-1 (PSGL1) from engineered CHO cells. For simplicity only the defined Lewis epitopes of the native protein *O*-glycome are shown. The enzymatic activity is monitored by the loss of Western blot signal originating from specific Le-epitope antibodies after enzyme incubation. The enzymes that display activity are highlighted by a pink box. The data are from a single (n=1) experiment.

**a**

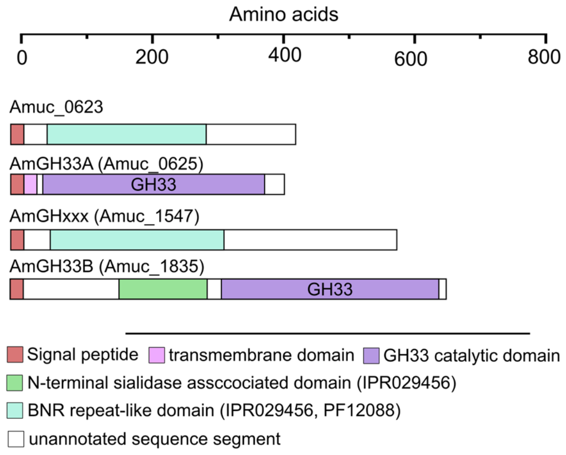


**b**

Amino acid sequence identity

| | Amuc_0623 | *Am*GH33A | AmGHxxx | *Am*GH33B |
|---|---|---|---|---|
| Amuc_0623 | 100 | 22.15 | 21.66 | 24.62 |
| *Am*GH33A | 22.15 | 100 | 26.89 | 28.75 |
| AmGHxxx | 21.66 | 26.89 | 100 | 22.98 |
| *Am*GH33B | 24.62 | 28.75 | 22.98 | 100 |

GH33 catalytic domain
BNR repeat-like domain (IPR029456, PF12088)

**c**

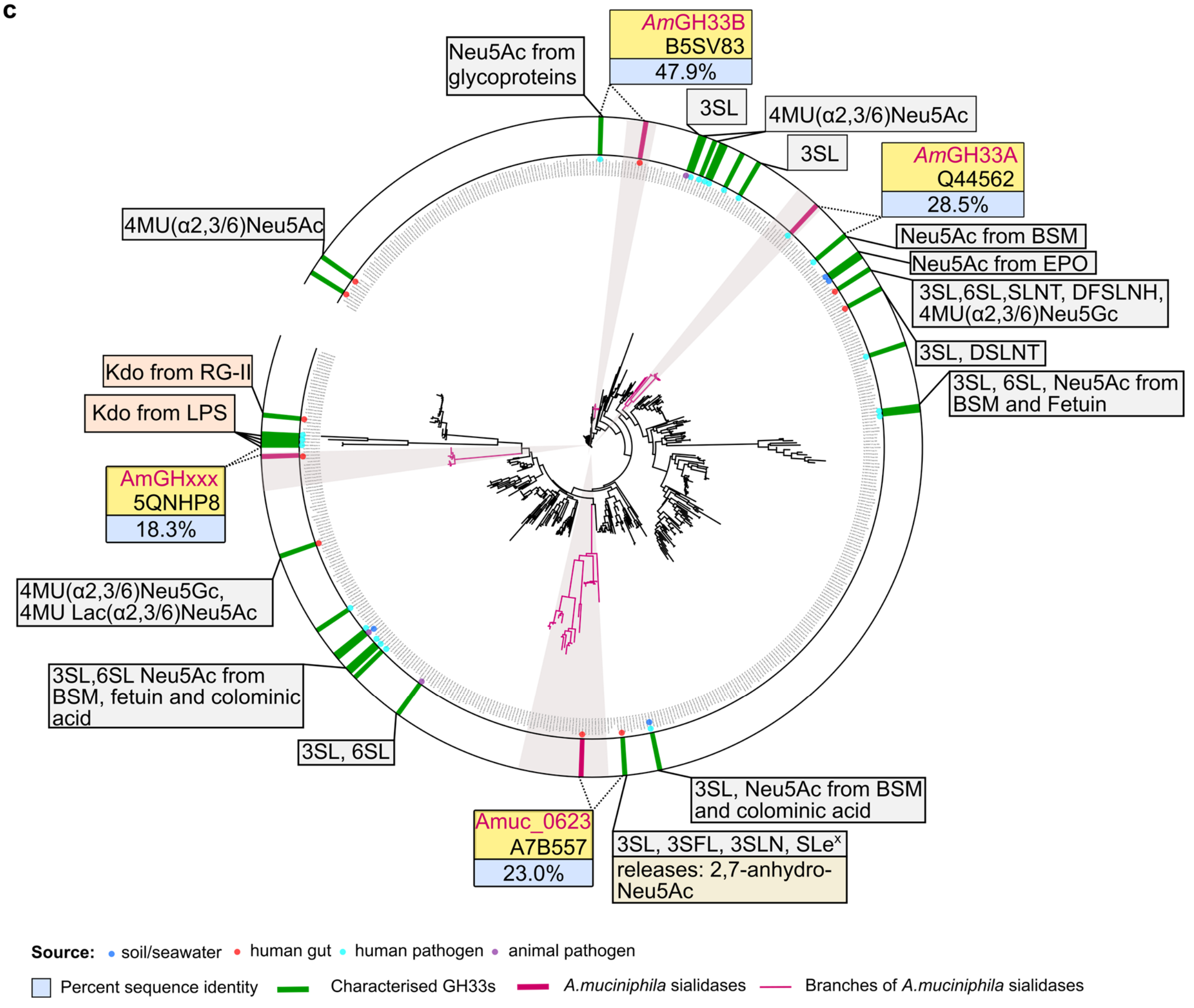


**Supplementary Fig. 7: Modular organization and phylogeny of *A. muciniphila* sialidases. a,** Modular organization of *A. muciniphila* assigned GH33 sialidases and distant orthologues with BNR-like domains, considered a sialidase signature. **b,** Amino acid sequence identity matrix for the sequences in a**.** **c,** Phylogenetic tree of 534 putative sialidases sequences (GH33 enzymes and BNR-like domain enzymes from the sequences in a and b). *A. muciniphila* sialidases and their closest characterized orthologues (indicated with their UniProt IDs) are in yellow boxes and the amino acid sequence identities between their catalytic modules are in the blue box. The source niches of the described sialidases are indicated by coloured circles and the substrates the enzymes have been shown to be active on are shown. The branches populated by the *A. muciniphila* enzymes are in pink and highlighted by a grey shadow. Characterized GH33 enzymes previously shown to release 2,7-anhydro Neu5Ac are highlighted by a dark yellow box and GH33 enzymes demonstrated to be active on 2-keto-3-deoxy-D-manno-octulosonic acid (Kdo) are highlighted by an orange box.

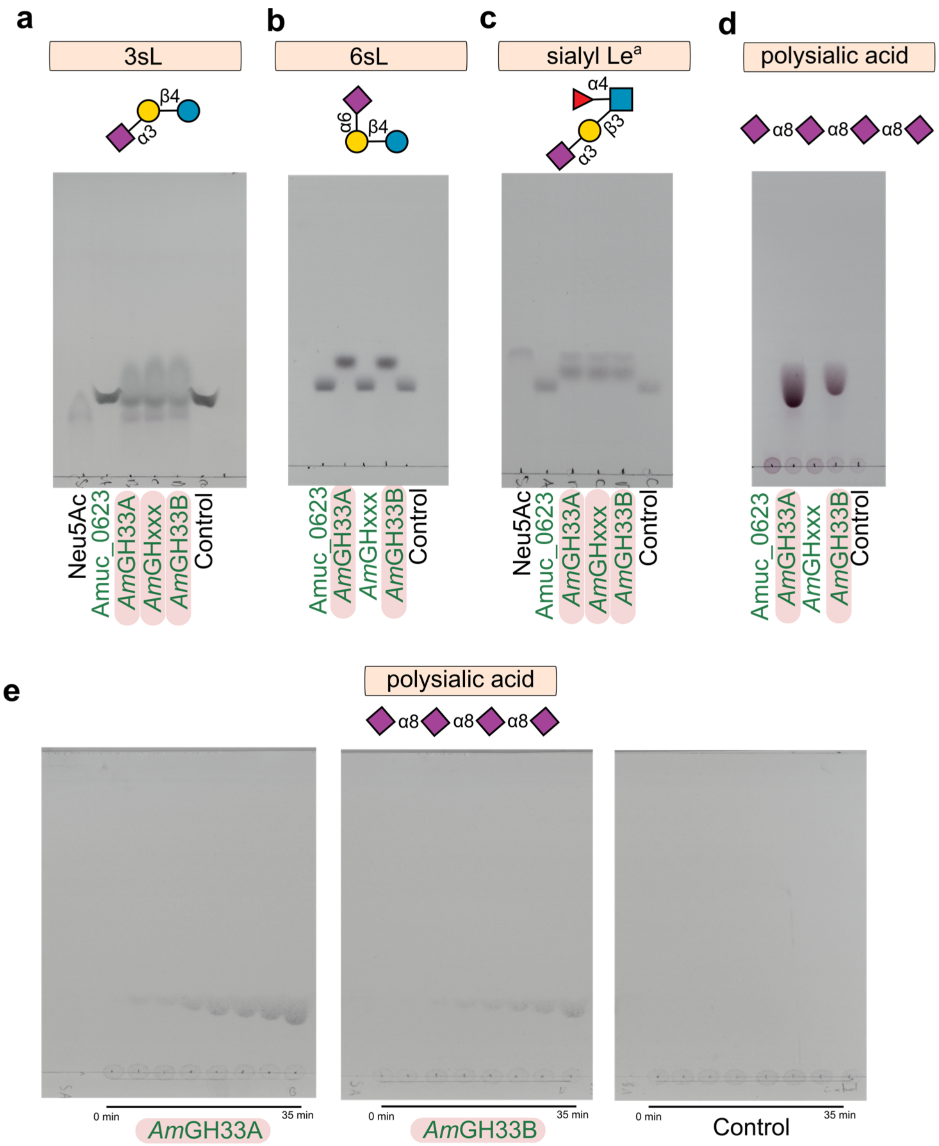


**Supplementary Fig. 8: Sialidase activity on oligosaccharides. a-e,** The sialidase activity on sialyl-substituted oligosaccharides and polysialic acid (Colominic acid) monitored by TLC. Active enzymes are highlighted by pink boxes. The data are from three independent experiments (n=3), whereby all analyses yielded similar results.

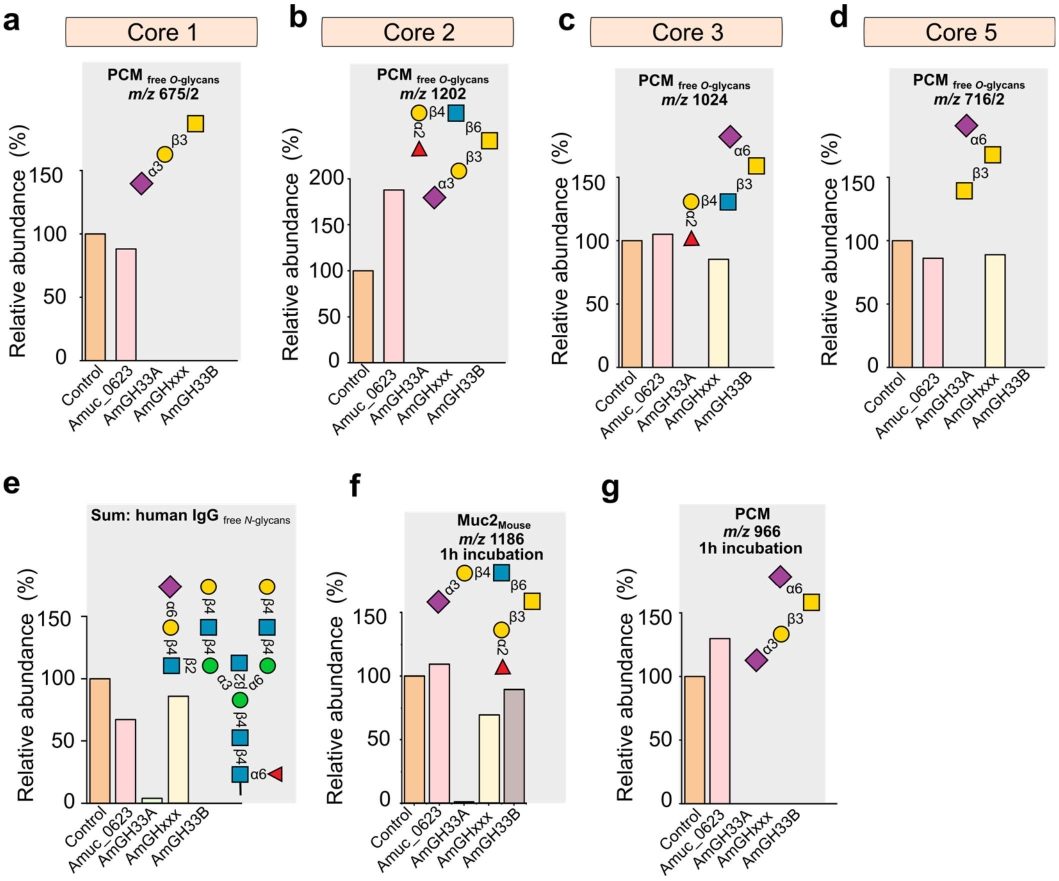


**Supplementary Fig. 9: Sialidase activity on mucin sialylated *O*-glycan motifs as well as immunoglobulin G *N*-glycans**. The subscripted "free *O*-glycans" indicated that the analyses were performed on released glycans, whereas absence of this subscript refers to analyses conducted on intact mucin **a-d,** The activity on different *O*-glycan cores from porcine colonic mucin (PCM). **e,** The sialidase activity on *N*-glycans from human IgG. **f,** Activity profiles showing equal activity of *Am*GH33A on the 6SGalNac epitope from PCM and Muc2$_{Mouse}$ (Muc2 from mouse) after 1 h incubation, while the activity of *Am*GH33B on the same epitope is markedly lowered on intact Muc2$_{Mouse}$. **g,** Activity profile showing the lowered activity of *Am*GH33B on the Sd$^{a}$ epitope from Muc2$_{Mouse}$ as compared to the activity of the enzyme on the same epitope from PCM after 1 h incubation. For isobaric glycans structures, the number after the slash (/n) denotes the corresponding structure in the LC-MS data file (see extended data). The data are from a single (n=1) experiment.

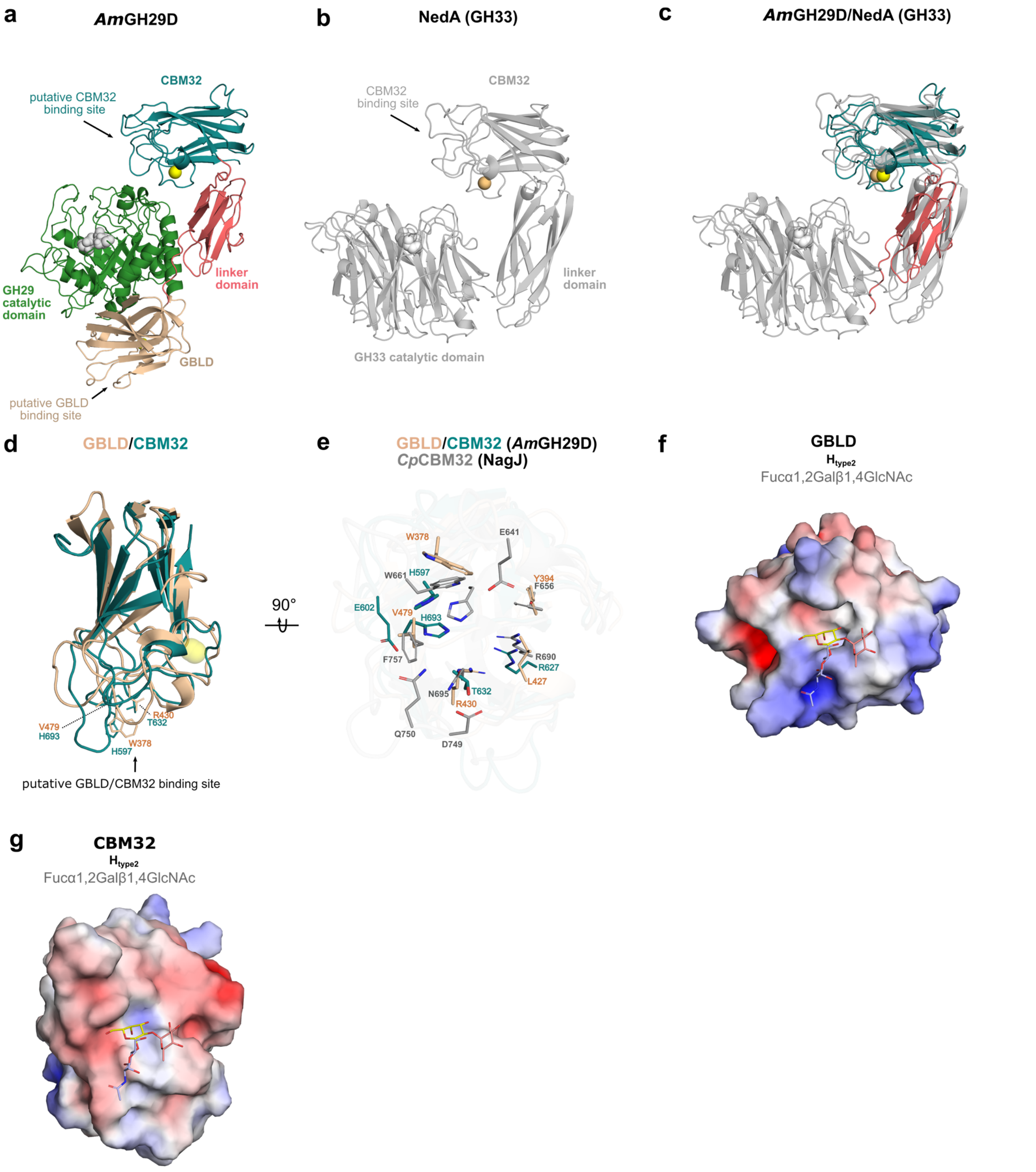


**Supplementary Fig. 10: Cobra strike pose architecture and putative binding domains of *Am*GH29D. a,** Overall structure of *Am*GH29D consisting of canonical catalytic N-terminal $(\beta/\alpha)_8$ domain (green), a β-sandwich barrel forming the galactose binding like domain (GBLD) (wheat coloured), a linker domain (salmon red) and a putative CBM32 (dark cyan) with a $Ca^{2+}$ (yellow sphere) binding site. The inferred catalytic nucleophile D190 and the acid/base E246 are shown as spheres (white). **b,** Overall structure of the *Micromonospora viridifaciens* GH33 sialidase (NedA, PDB: 1WCQ) that displays a similar "Cobra strike pose" architecture as *Am*GH29D. The catalytic residues are highlighted as spheres (grey) and the bound $Na^+$ is shown as orange sphere, **c,** Structural alignment (Dali server) of NedA with the linker and CBM32 domain of *Am*GH29D showing a similar juxtapositioning above the active site (here dubbed as a Cobra strike pose). **d,** Structural alignment of the GBLD and the CBM32 domains of *Am*GH29D with the bound $Ca^{2+}$ (yellow) and with residues putatively involved in ligand binding represented as sticks. **e,** Comparison of the ligand-binding sites of the biochemically characterized CBM32 from *Clostridium perfringens* (*Cp*CBM32, PDB: 2J7M) with GBLD and the CBM32 from *Am*GH29D. The analysis shows that the aromatic stacking tryptophan in the previously characterized CBM32 is shared with GLBD, which otherwise possesses more apolar residues in the potential binding site as compared to both CBM32s. **f,** Electrostatic surface representation of the GLBD (generated using the APBS plugin in Pymol) illustrating the apolar surface of the putative binding site. **g,** Electrostatic surface representation of the putative CBM32 showing a more polar putative binding site as compared to the GLBD (g). (**f** and **g)** The H2 antigen trisaccharide bound in *Cp*CBM32 with fucose shown in pink, *N*-acetylglucosamine in blue and galactose in yellow is also shown in the GBLD and CBM32 from *Am*GH29D after structural alignment with the *Cp*CBM32 (2J7M) to visualize the putative binding site area.

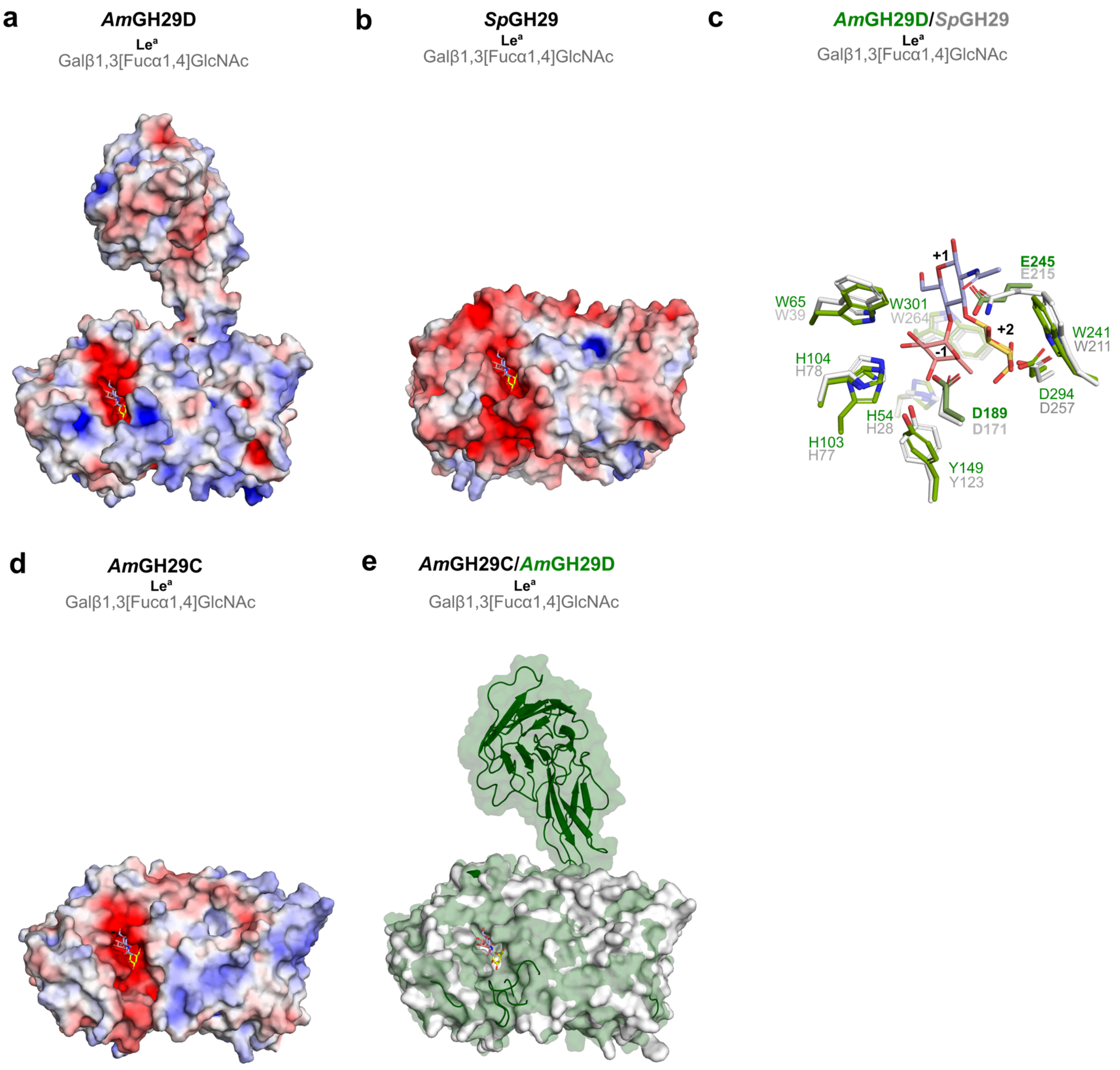


**Supplementary Fig. 11: Comparison of the ALPHA fold model of *Am*GH29C and the crystal structure of *Am*GH29D. a,** Surface representation of *Am*GH29D (colored according to electrostatic potential APBS plugin in Pymol) showing the negatively charged active site surrounded by positively charged patches. **b,** Electrostatic surface representation of a GH29 from *Streptococcus pneumonia* (*Sp*GH29, 6OR4; closest structural orthologue of *Am*GH29D) showing a largely negatively charge patch flanking the active site as opposed to *Am*GH29D. **c,** Stick representation (green) of the *Am*GH29D active site shows a GH29 typical highly -1 subsite architecture and ligand recognition when compared to *Sp*GH29 (white, stick representation). The catalytic residues from *Am*GH29D (D190 and E246) are highlighted by bold labels. **d,** Electrostatic surface representation of the ALPHA fold model of the *Am*GH29C catalytic domain. **e,** Superimposition of the *Am*GH29C catalytic domain ALPHA fold model (white solid surface) with *Am*GH29D (green semi-transparent surface) showing large loops in *Am*GH29D restricting the active site, potentially hindering the accommodation of larger complex and/or heavily substituted fuco-*O*-glycans or internally fucosyl substituted glycans as opposed to *Am*GH29C.

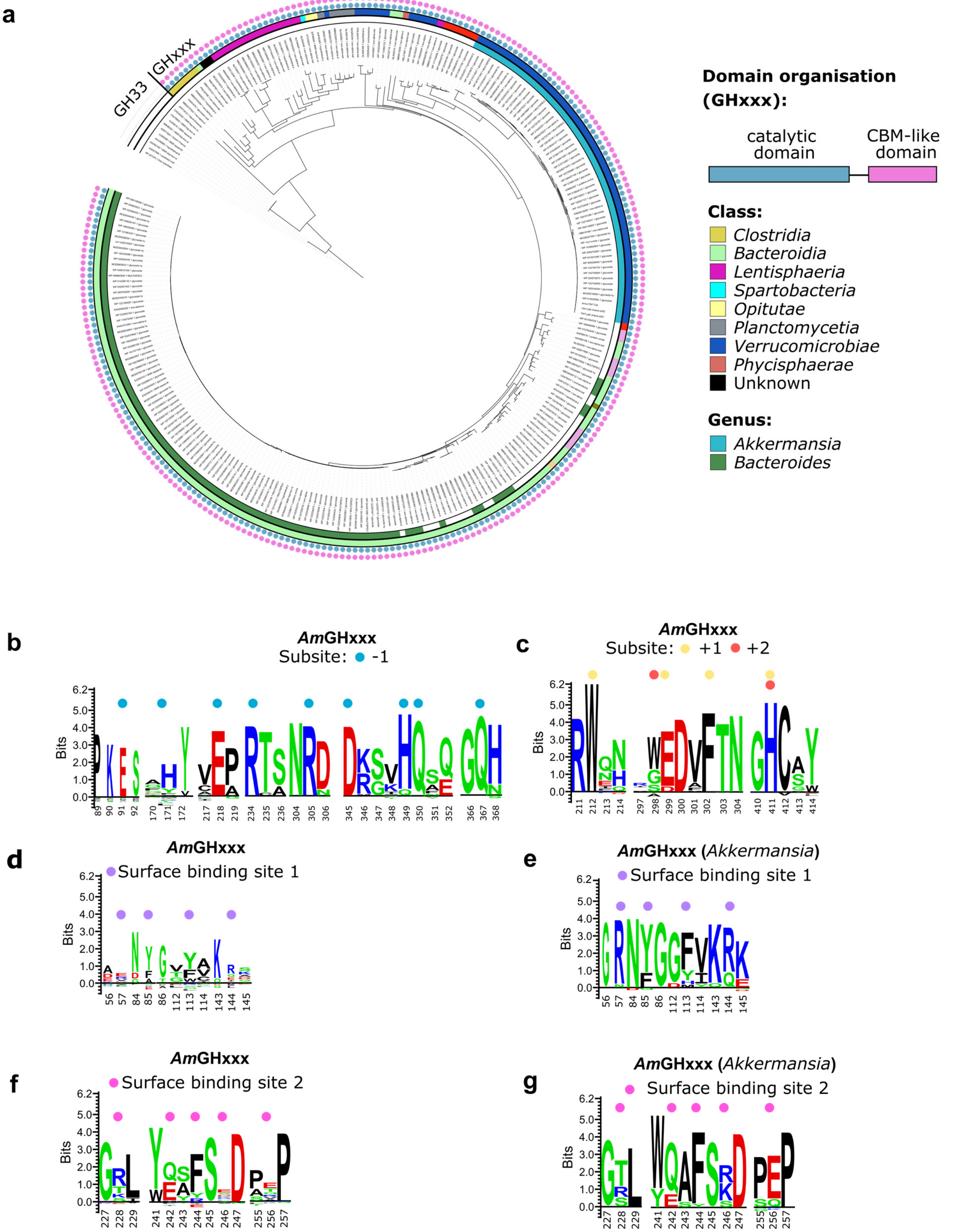


**Supplementary Fig. 12: Phylogenetic tree of GHxxx as well as active site and surface binding site motif conservation. a,** Phylogenetic tree of 335 GHxxx sequences generated using *Am*GHxxx as a query (see Materials and methods section). The modular architecture of the catalytic module and the CBM-like domain is conserved throughout the family and the taxonomic class and genus affiliations of the sequences is shown. **b,** Sequence logo of subsites +1 and +2 subsites. **c,** Sequence logo of subsite -1. **d,** Sequence logo of the putative surface binding site 1 residues across GHxxx. **e,** same as **d**, but including only GHxxx sequences from the *Akkermansia* genus. **f,** Sequence logo of the putative surface binding site 2 across GHxxx. **g,** same as **f**, but including sequences only from the *Akkermansia* genus.

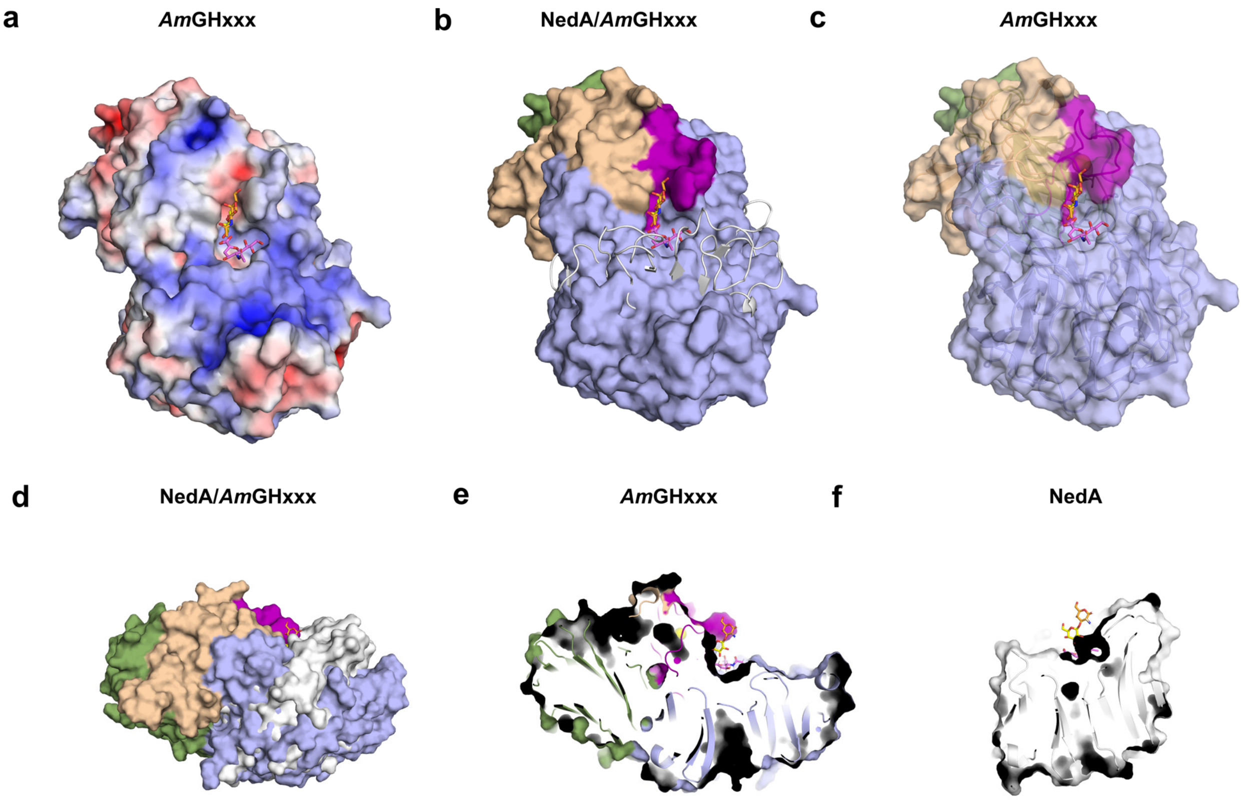


**Supplementary Fig. 13: Architecture of *Am*GHxxx and comparison to GH33 sialidases. a,.** Electrostatic surface representation of *Am*GHxxx showing large positively charged patches surrounding the catalytic site. **b,** Superimposition of *Am*GHxxx with the closest structurally characterized orthologue, the GH33 sialidase NedA from *Micromonospora vifidifaciens* (1EUS), showing that the elongated loops that join the strands of the β-propeller in GH33 (gray cartoon) are markedly shortened in *Am*GHxxx, resulting in open side of the active site as opposed to NedA. By contrast, longer loops together with the $Ca^{2+}$ binding and the B domains pack onto the β-propeller forming the binding site for the T-antigen disaccharide moiety, which lacks in GH33 enzymes. *Am*GHxxx is shown as a surface colored according to the domains: the catalytic domain in light blue, the $Ca^{2+}$ (yellow sphere) binding domain in violet, the B domain in wheat and the C-terminal β-sandwich domain in green. **c,** Semi-transparent surface representation of *Am*GHxxx colored as in b showing that the active site of *Am*GHxxx is shaped by the $Ca^{2+}$ bindings site, the B domain and two large loops from the catalytic domain. **d,** Superimposition of *Am*GHxxx (solid surface, colored as in **b**) with NedA (solid surface, white) showing differences in active site architecture. **e,** Same view of *Am*GHxxx as d but represented as carved solid surface (coloured as in **b**) highlighting the "sun-chair" architecture of the *Am*GHxxx active site. **f,** Same view of NedA as in d but represented as carved solid surface (white) showing the flat surface NedA potentially lacking the aglycone (+) ligand binding sites. Structural alignments of *Am*GHxxx and NedA (panels **b** and **d**) were performed using the DALI server.

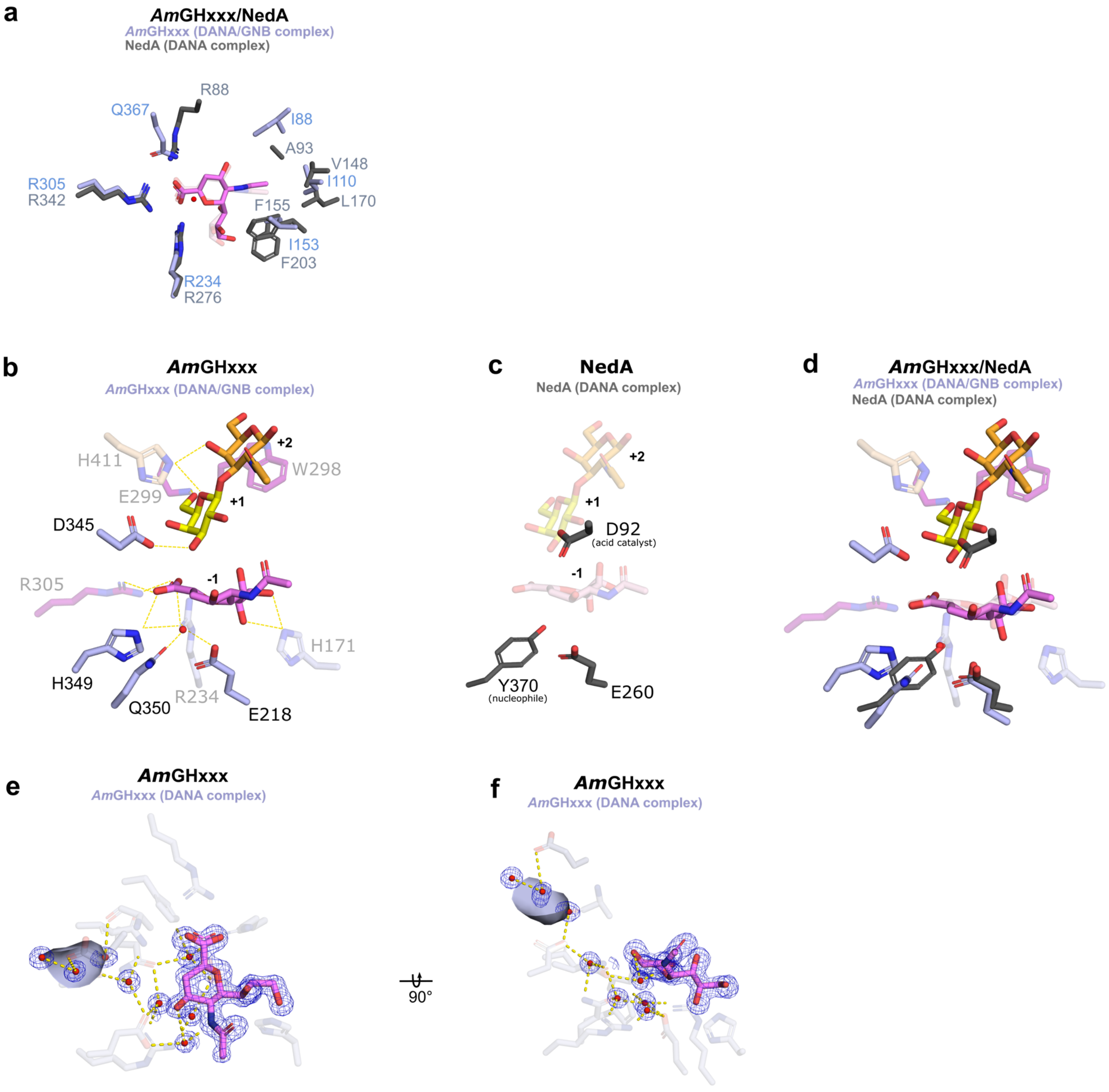


**Supplementary Fig. 14 Catalytic site signatures of *Am*GHxxx as compared to the closest GH33 sialidase. a,** Superimposition of the catalytic sites of *Am*GHxxx (light blue stick representation, the DANA inhibitor in dark pink, Gal in yellow, GalNAc in dark yellow and an ordered water molecule as red sphere) and NedA, the GH33 from *Micromonospora vifidifaciens* (1EUS, grey stick representation, the DANA inhibitor in a semi-transparent and light pink). The top view shows that two of the three arginine residues of the conserved R triad in GH33, are also conserved in *Am*GHxxx, whereas a glutamine substitutes the third arginine. Residues with similar chemistry flank the *N*-acetyl group of the DANA inhibitor, while two phenylalanines, one of which packs onto the DANA inhibitor in the catalytic site, are lacking in *Am*GHxxx **b,** The catalytic site of *Am*GHxxx showing the recognition of the DANA inhibitor and T-antigen disaccharide (GNB) (stick representation with the amino acid residues coloured according to domain, similar to Supplementary Fig. 11b). **c,** The catalytic machinery of the GH33 NedA (stick representation) showing the catalytic tyrosine nucleophile, an adjacent conserved glutamate and the catalytic acid. The DANA and GNB from *Am*GHxxx are also visualized to keep the perspective of the active site. **d,** Superimposition of the *Am*GHxxx catalytic site (same colouring scheme as b) and NedA (grey sticks, DANA in semi-transparent and light pink) showing the substitution of the catalytic nucleophile in GH33 to a glutamine (Q350) in *Am*GHxxx which is preceded by a histidine (H349). These two residues as well as an invariant glutamate (E218) and one of the conserved arginine (R234) are potentially hydrogen bonded to a likely catalytic water molecule (see b) that overlays perfectly with the oxygen in the catalytic tyrosine in GH33. Thus, the catalytic water is positioned for nucleophilic attack at the C2 of the sialyl (or inhibitor) unit instead of the tyrosine in GH33. An invariant aspartate (D345) in *Am*GHxxx, which is hydrogen bonded to the C3-OH group of the bound galactosyl unit at subsite +1 (see b) potentially act as catalytic acid as opposed to the aspartate in GH33 (D92) which act as general acid/base. **e,** The $F_o - F_c$ electron density maps (blue mesh) of the DANA inhibitor and a solvent tunnel connecting the bulk of the solvent and the catalytic site. **f,** same representation as in e but rotated 90 degrees along the x-axis.

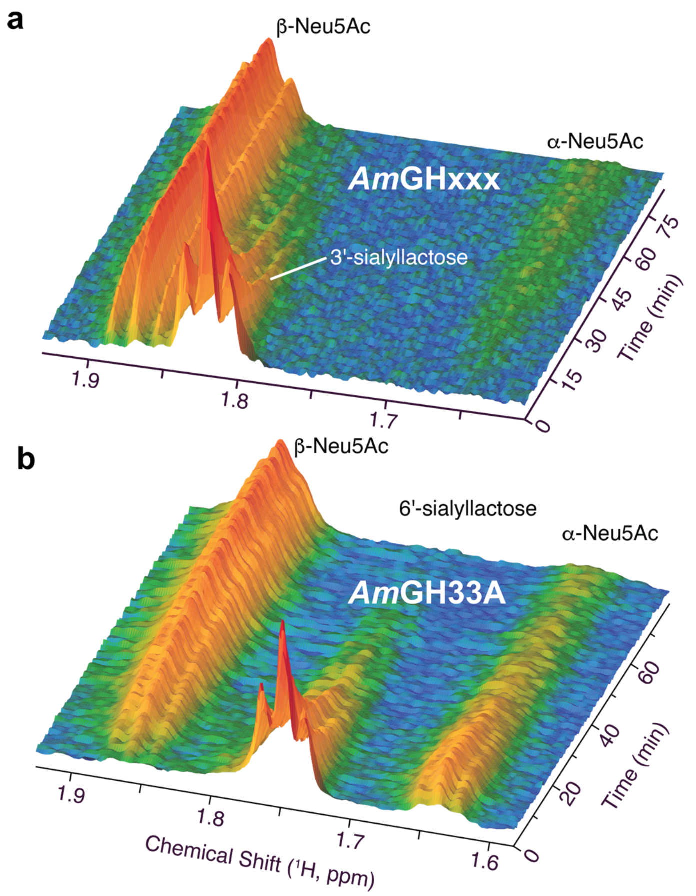


**Supplementary Fig. 15: The NMR analysis of the inverting mechanism of *Am*GHxxx. a,** Time series of $^{1}$H NMR real time spectra of the *Am*GHxxx-catalysed conversion of 3´-sialyllactose at 310 K and pH 6.8. The spectral region containing the axial $^{1}$H-3 signal in sialic acid (Neu5Ac) is depicted. The spectral series showed that β-Neu5Ac is the initial product of the reaction. Some α-Neu5Ac emerges as the reaction progresses due to mutarotation. These data provide evidence that the hydrolysis of Neu5Ac proceeds with the inversion of anomeric configuration in the previously undescribed GH174 family. **b,** Control experiment that shows a time series of $^{1}$H NMR real time spectra of the *Am*GH33B catalysed conversion of 6´-sialyllactose at 310 K and pH 6.8. The spectral region containing the axial $^{1}$H-3 signal in sialic acid (Neu5Ac) is depicted. The spectral series showed that α-Neu5Ac is the initial product of the reaction that mutarotates to β-Neu5Ac with a half time of about 80 min to reach about 90% of the total Neu5Ac in the solution. These data are consistent with the known retaining mechanism within GH33. The data are from a single (n=1) experiment.

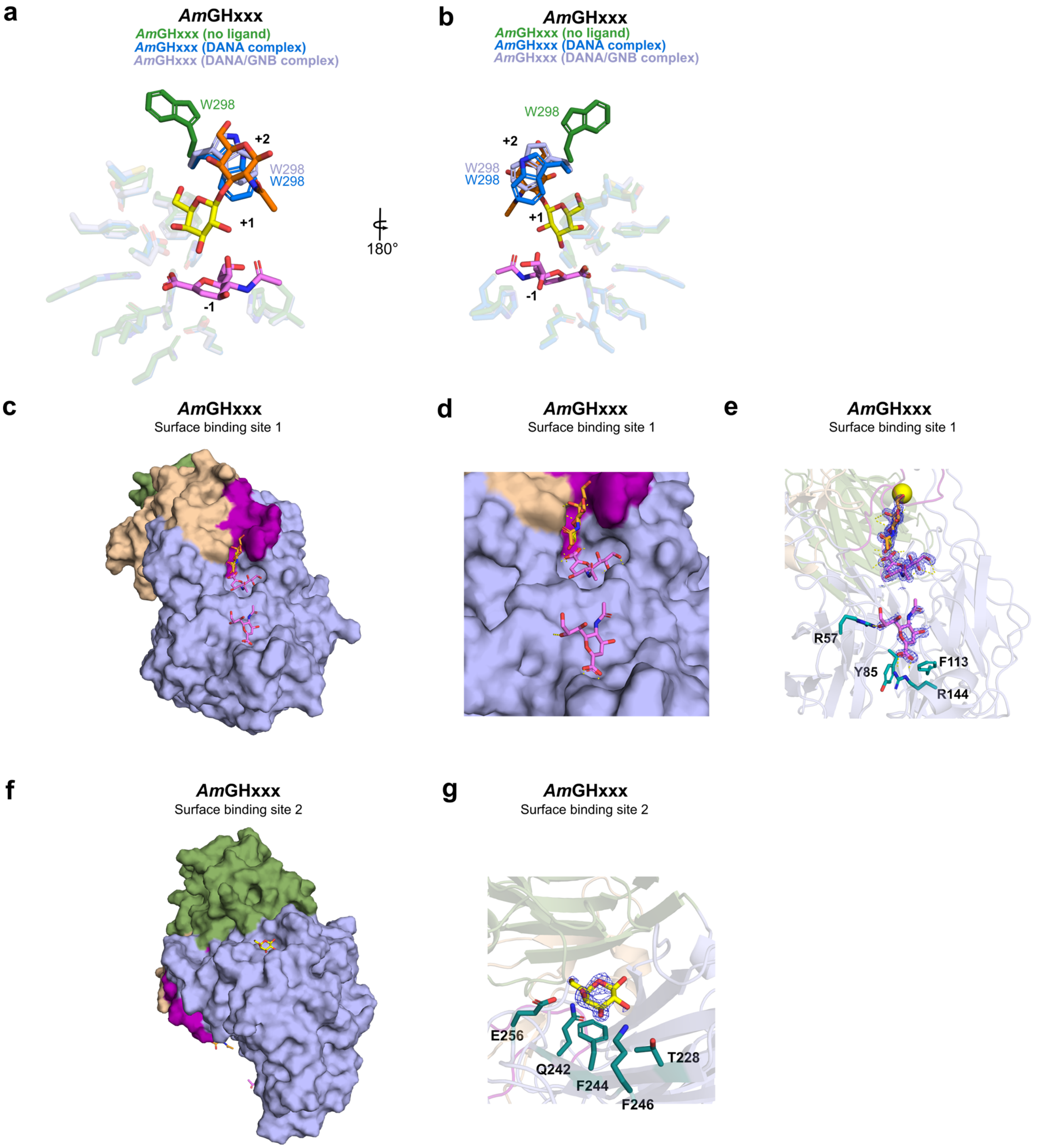


**Supplementary Fig. 16: Ligand binding at the active site and secondary surface binding sites of *Am*GHxxx. a,** Comparison of the ligand free (green sticks), DANA bound (blue) or DANA+GNB bound (light purple) structure, showing an induced fit flipping movement of a tryptophan sidechain to provide aromatic stacking for the GalNAc at subsite +2 in the two ligand-bound structures. **b,** The same as **a**, but turned 180°. **c,** Surface binding site 1 adjacent to the active site with a DANA molecule bound. **d,** Zoom in view as in **c**. **e,** potential binding residues at 4 Å distance from the modelled DANA molecule. **f,** The surface binding site 2 at the opposite side of the active site with a modelled galactose unit bound at a shallow groove. **g,** The potential binding resides of the modelled Gal with a phenylalanine aromatic stacking interaction flanked by polar residues. (e and g) Unbiased $F_o - F_c$ electron density maps are represented as blue mesh.

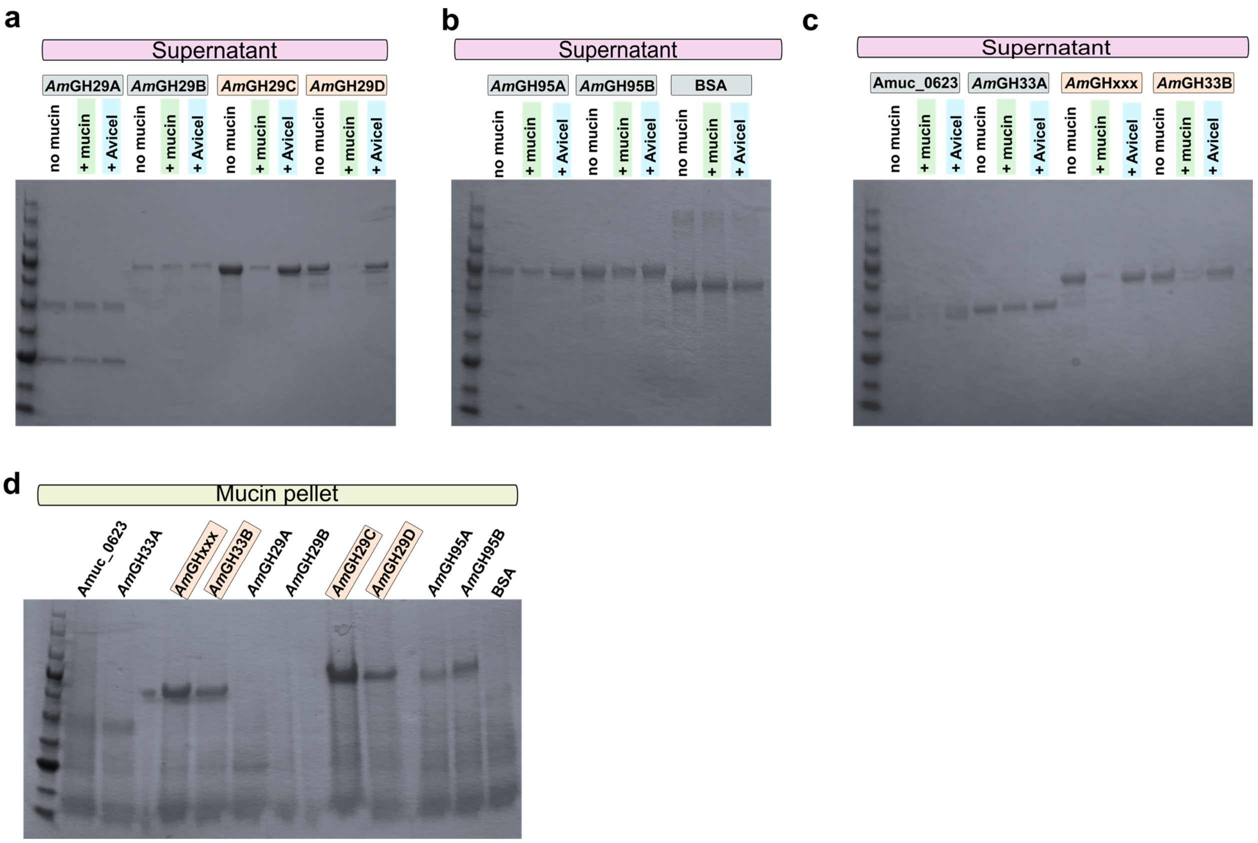


**Supplementary Fig. 17: Binding of *A. muciniphila* fucosidases and sialidases to mucin. a-d,** Representative SDS-PAGE gels showing pulldown binding assays of *A. muciniphila* fucosidases, sialidases and of a negative control protein (BSA) binding to insoluble PCM and Avicel. The data are from two independent experiments (n=2) whereby all analyses yielded similar results.

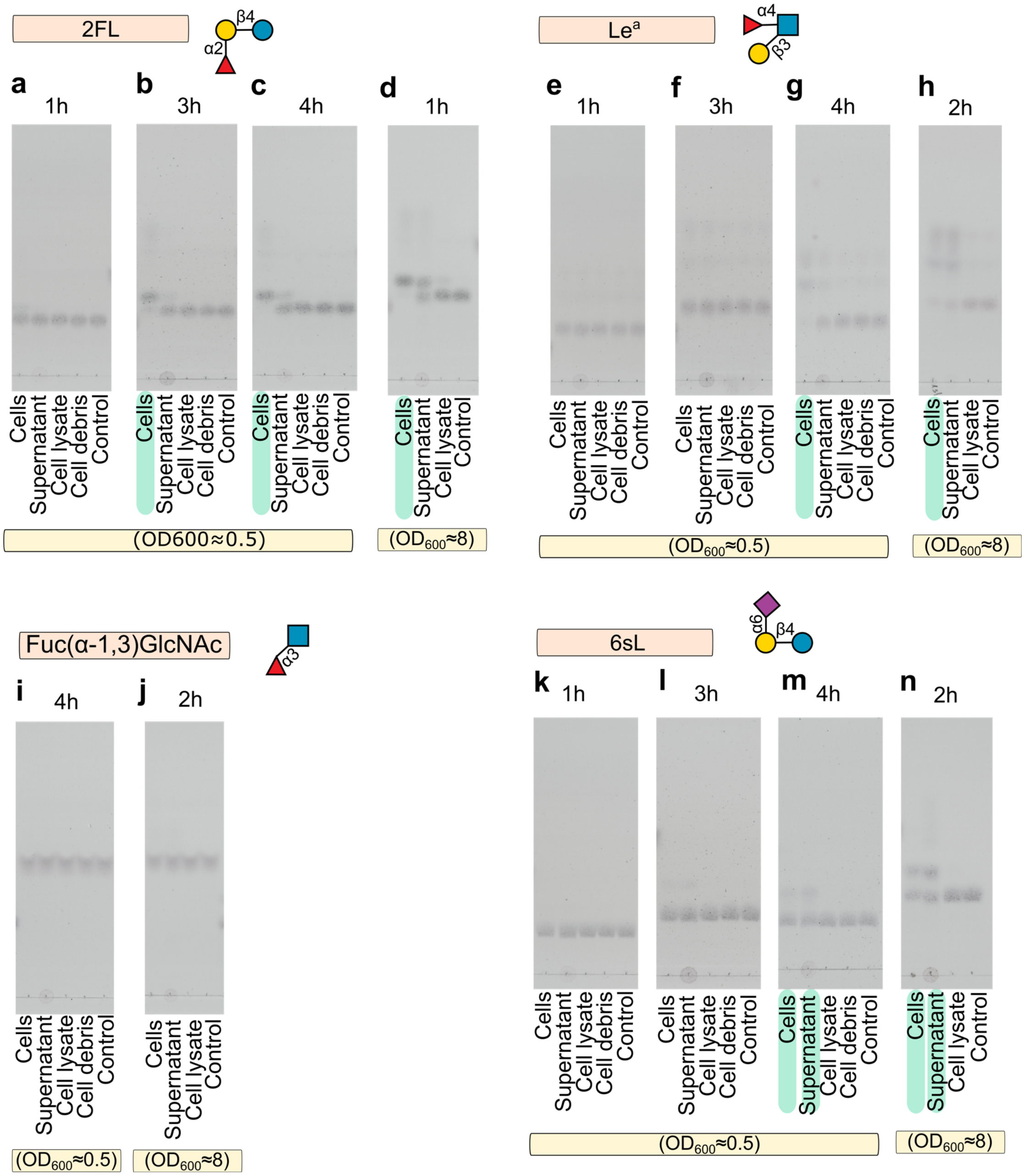


**Supplementary Fig. 18: Localization of fucosidase and sialidase activities encoded by *A. muciniphila*. a-d,** α-1,2-fucosidase activity of *A. muciniphila* as assayed on 2FL with intact cells, culture supernatants, cell lysates, cell debris and a buffer control. **e-h,** α-1,4-fucosidase activity of *A. muciniphila* as assayed on Le[a] trisaccharide on different culture fractions with a buffer as a negative control. **i-j**. α-1,4-fucosidase activity of *A. muciniphila* as assayed on Fuc α-1,3 GlcNAc using same culture fractions and control. **k-n,** α-2,6-fsialidase activity of *A. muciniphila* as assayed on 6SL with intact cells, culture supernatants, cell lysates, cell debris and a buffer control. The experiments have been performed on cells grown porcine gastric mucin. Fractions showing highest activity are highlighted with green boxes. The data are from three independent experiments (n=3) whereby all analyses yielded similar results.

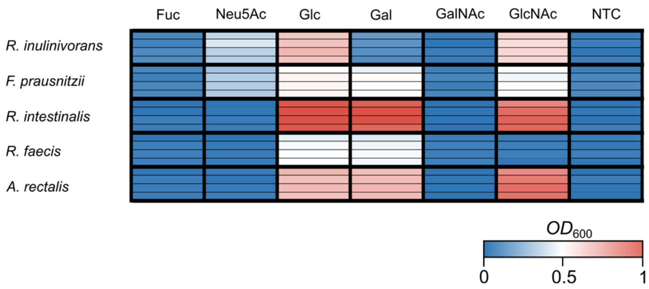


**Supplementary Figure 19. Growth of butyrate producing Lachnospiraceae on monosaccharides from mucin. a,** Growth of *Roseburia inulinivorans, Roseburia intestinalis*, *Roseburia faecis, Agathobacter rectalis, Faecalibacterium prausnitzii* on YCFA supplemented with 0.5 % (w/v) monosaccharides from mucin after 24h. Growth experiments were performed in four independent biological replicates (n=4).
